# PII signal transduction superfamily acts as a valve plug to control bicarbonate and ammonia homeostasis among different bacterial phyla

**DOI:** 10.1101/2023.08.10.552651

**Authors:** Michael Haffner, Wen-Tao Hou, Oliver Mantovani, Peter R. Walke, Ksenia Hauf, Marina Borisova, Martin Hagemann, Cong-Zhao Zhou, Karl Forchhammer, Khaled A. Selim

## Abstract

Life on Earth relies on carbon and nitrogen assimilation by RubisCO and GS-GOGAT enzymes, respectively, whose activities depend on a constant supply of inorganic carbon (C_i_) and nitrogen (N). Members of the PII signal transduction superfamily are among the most ancient and widespread cell signaling proteins in nature. One of their most highly conserved functions is controlling C_i_- and N-transporters, a feature found in different phyla of the Archaea and in both Gram-positive and Gram-negative bacteria. Recently, we identified the PII-like protein SbtB as C_i_-sensing module, mainly controlling the HCO_3_^-^ transporter SbtA in cyanobacteria. Similar to canonical PII proteins, SbtB is able to bind the adenine nucleotides ATP and ADP. Unlike those, it also binds AMP and preferentially binds the second messenger cAMP and c-di-AMP. The functional significance of the binding of different adenyl-nucleotides to SbtB has remained elusive, particularly in the context of the interaction of SbtB with SbtA. By a combination of structural, biochemical and physiological analysis, we revealed that by binding to SbtA, SbtB acts as unidirectional valve, preventing the reverse transport of intracellular enriched bicarbonate. This mechanistic principle holds true for the PII protein from *Bacillus* acting on the ammonium transporter AmtB, suggesting an evolutionary conserved role for PII superfamily proteins in controlling unidirectional flow of different transporters.

## Introduction

The PII signaling superfamily comprises the most ancient and widespread signal transduction proteins found in all domains of life (Forchhammer et al. 2022; Wheatley et al. 2016; Selim et al. 2018, 2021). The well-studied canonical PII proteins, which are involved in the regulation of carbon/nitrogen homeostasis, are characterized by a highly conserved trimeric structure with ferredoxin-like fold (Forchhammer & Selim 2020; Selim et al. 2020). Despite the similar 3D structures, PII-like proteins show restricted amino acid sequence conservation and evolved to fulfill diverse, yet not completely understood cellular functions.

Cyanobacteria are oxygenic phototrophs that assimilate CO_2_ into organic compounds using water and light energy. For efficient CO_2_ fixation, cyanobacteria evolved an efficient carbon concentrating mechanism (CCM) that employs high affinity inorganic carbon (C_i_; referring to CO_2_ and HCO_3_^-^) uptake systems to augment intracellular C_i_ levels (Forchhammer & Selim 2020; Hagemann et al. 2021). The sodium-dependent bicarbonate transporter SbtA is an important component of the cyanobacterial CCM and becomes highly expressed under C_i_ limitation from the *sbtA-sbtB* operon, which also encodes the conserved PII-like protein SbtB (Mantovani et al. 2022). Similar to canonical PII proteins, SbtB is able to bind the adenine nucleotides ATP and ADP, but unlike those, it also binds AMP. Moreover, SbtB preferentially binds the second messengers cAMP (Selim et al. 2018), which is known as carbon-status indicator (Chen et al. 2000; Steegborn et al. 2005), and c-di-AMP, which is involved in global cellular homeostasis (Selim et al. 2021; Mantovani et al. 2022, 2023).

Detailed structural characterization of various cyanobacterial SbtB homologues showed that they display typical PII-protein characteristics, being homo-trimeric proteins with three characteristic loop regions (T-, B- and C-loops) (Selim et al. 2018, 2021, 2023; Kaczmarski et al. 2019). These loop regions are located near the inter-subunit clefts and play a major role in ligand binding and intramolecular signaling. Intriguingly, the structures of *Synechocystis* SbtB:AMP (PDB: 5O3R) and SbtB:cAMP (PDB: 5ORQ) complexes did not reveal any conformational changes on the flexible surface-exposed T-loop, the protein interaction module of PII superfamily proteins that usually responds to effector molecule binding. Another striking feature of many SbtB proteins, including *Synechocystis* SbtB (hereafter *Sc*SbtB), is a C-terminal extension (referred to as R-loop) with a conserved C_105_GPxGC_110_ motif to which we ascribed a redox-regulatory function due to a disulfide bond formation between C_105_ and C_110_ (Selim et al. 2023).

In summary, we revealed that *Sc*SbtB acts as C_i_-sensor module by integrating the energy/redox-state of the cell and second messengers cAMP and c-di-AMP, and thereby it plays a central role in the regulation of the entire CCM, similar to the role of the canonical PII protein in controlling carbon/nitrogen balance in central metabolism (Forchhammer et al. 2022; Selim et al. 2018, 2021; Mantovani et al. 2022). While the evolutionary conserved role of cAMP/AMP as indicators of cellular carbon status is well understood (Steegborn et al. 2005; Selim et al. 2018; Fang et al. 2021), the specific function of ATP/ADP adenyl-nucleotide binding to SbtB has remained unresolved. Several studies assigned SbtB to the regulation of SbtA through direct protein-protein interaction (Du et al. 2014; Selim et al. 2018, 2023; Liu et al. 2021), in analogy to the canonical PII member GlnK-mediated inhibition of the ammonium transporter AmtB in Gram-negative bacteria under nitrogen excess conditions to prevent an over-accumulation of ammonium (Conroy et al. 2007; Gruswitz et al. 2007; Boogerd et al. 2011; Forchhammer et al. 2022). Recently, we showed that SbtB displays an unusual slow ATP/ADP apyrase (diphosphohydrolase) activity that is controlled by the C-terminal R-loop in response to the redox-state of the cell (Selim et al. 2023). The SbtB apyrase activity leads to a conformational change in the surface-exposed T-loop, hence the ATP/ADP binding could act as a molecular switch that drives a conformational change in the T-loops. Although its physiological role has remained obscure, the redox-regulated apyrase activity of SbtB may link the photosynthetic electron transport to the regulation of SbtA mediated bicarbonate transport.

Recently, we and others solved the structures of SbtA and SbtA:SbtB complex (Fang et al. 2021; Liu et al. 2021). We showed that SbtA is a secondary-active transporter and operates by an elevator-type mechanism, with alternating outward and inward open substrate transport sites (Liu et al. 2021). Furthermore, these studies revealed symmetric binding of the trimeric SbtB to a trimeric SbtA. In the SbtA:SbtB complex, the SbtB T-loops lock SbtA in the inward-facing state, thereby assumed to inhibit the SbtA bicarbonate transporter activity (Fang et al. 2021). This strongly implies that SbtB exerts a direct regulatory function on SbtA in response to the metabolic state of the cell through binding to various adenine nucleotides, which may induce conformational changes in the SbtA-interacting T-loop in analogy to canonical PII proteins (Forchhammer & Selim 2020; Forchhammer et al. 2022). In fact, it was demonstrated that the conformation of the T-loop in the SbtA:SbtB complex is favored by AMP binding but is sterically incompatible with cAMP binding (Selim et al. 2018, 2023; Fang et al. 2021). The conclusion of an inhibitory SbtA:SbtB complex is, however, in conflict with the fact that the complex forms when high bicarbonate uptake activity is required. It also conflicts with the phenotype of Δ*sbtB* deficient mutants, which showed growth impairment under low C_i_ supply (Selim et al. 2018), suggesting that SbtB is indeed required for efficient C_i_ uptake.

Deeper insights in the mechanism, how SbtB affects the SbtA transporter can be obtained by comparing different snapshots from recent structural studies (Fang et al., 2021; Liu et al. 2021). In absence of AMP, SbtB is still able to interact with SbtA in the cytoplasmic inward-facing state, although the SbtB T-loop was disordered (PDB: 7EGL). In this the structure, the HCO_3_^−^ substrate was clearly defined toward the cytoplasm (Fang et al. 2021). We also revealed that the SbtB in the AMP state induces the cytoplasmic inward-open conformation of SbtA by separating the SbtA core-domain from the gate-domain, where the structured SbtB T-loop only partially locks the HCO_3_^-^ exit tunnel of SbtA, leaving SbtA in an inward-open conformation (PBD: 7CYF) (Liu et al. 2021). This implies that SbtB within the SbtA:SbtB complex remains in dynamic to regulate the SbtA transport activity in response to the concentrations of intracellular adenyl-nucleotides and the redox-state of the cell. Another important aspect, so far not considered, is the fact that the transport mechanism of the secondary-active transporter to which SbtA belongs, is reversible (Sauer et al. 2022).

This work aimed to gain a deeper understanding of the mechanism of how SbtB affects transport activity of SbtA in *Synechocystis*. We show that formation of the SbtA:SbtB complex doesn’t inhibit the HCO_3_^-^ uptake but rather blocks leakage of bicarbonate in response to the energy state of the cell, suggesting a novel function of PII-like proteins as unidirectional valve in transport complexes preventing reverse transport of the substrate along the concentration gradient to the outside. We also expanded our model by reexamining the effect of the canonical PII protein in *Bacillus subtilis*, GlnK, on ammonium transport. The paradigm of a purely inhibitory action of PII-proteins on transporters, as derived from studies on GlnK-AmtB interaction in Gram-negative bacteria (Gruswitz et al. 2007), conflicts also with the finding that the homologue GlnK:AmtB complex in *B*. *subtilis* is strongly expressed under conditions of nitrogen-limitation, where active ammonium uptake is required (Detsch & Stülke 2003; Heinrich et al. 2006; Kayumov et al. 2011). Here we show that *B. subtilis* GlnK also acts as a valve to prevent ammonium leakage through AmtB, suggesting that this mechanism is a widely occurring feature among members of the PII superfamily.

## Results

### SbtB nucleotides diphosphohydrolase (apyrase) activity requires T-loop arginines and an oxidized R-loop

In our previous structural characterization of *Sc*SbtB (Selim et al. 2023), we observed that SbtB possesses a novel type of diphosphohydrolase (apyrase) activity, which is redox-dependent, converting either ATP or ADP to AMP. In summary, in the oxidized SbtB state, when the R-loop is folded via the C_105_-C_110_ disulfide bridge (PDBs: 7R2Y, 7R2Z and 7R30), the adenine nucleotides (ATP or ADP) are prone to hydrolysis due to a steric conflict of the oxidized R-loop with the T-loop, which forces the β- and γ-phosphates of ATP/ADP into a highly strained conformation, exposing the phosphates to hydrolytic attack. By contrast, *Sc*SbtB in the reduced state displays an unfolded R-loop, which allows the T-loop to adopt an ATP/ADP binding conformation, in which these nucleotides are protected from hydrolysis (PDBs: 7R31 and 7R32). Moreover, the T-loop arginine residues R_43_ and R_46_ were shown to be important for both ADP and ATP binding, as they coordinate the β- and γ-phosphates and K_40_ is situated close to the metal ion coordinating the β- and γ-phosphates (Selim et al. 2023). To experimentally validate the function of those residues, we created variants of SbtB, in which the T-loop residues K_40_, R_43_ and R_46_ were substituted to alanine. Using isothermal titration colorimetry (ITC), we tested the ability of the protein variants to bind adenyl nucleotides (cAMP, AMP, ADP and ATP). In comparison to wildtype *Sc*SbtB, all the T-loop variants were strongly impaired in ADP and ATP binding as indicated by high *K_d_* values, but less impaired in AMP or cAMP binding (Fig. 1A, Fig. S1, and Table 1), a property that was most prominently seen in the SbtB-R_46_A variant.

**Fig. 1:**
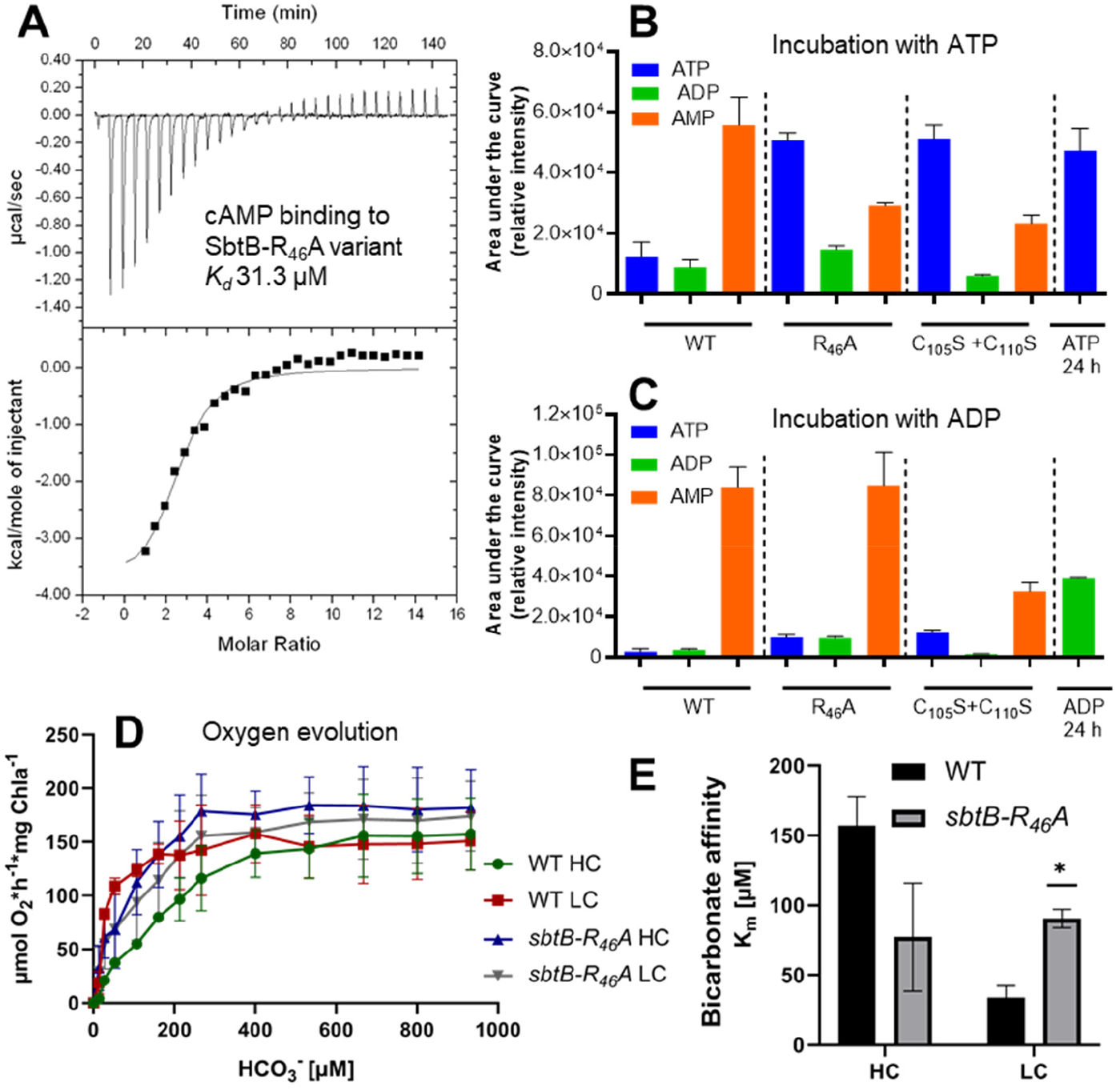
Characterization of SbtB T- and R-loop variants. (A) Isothermal titration calorimetry (ITC) shows that SbtB-R_46_A variant can still bind cAMP efficiently. (B,C) Nucleotide hydrolysis by SbtB wildtype and its variants (R_46_A and C105S+C110S [+ve control]) were incubated with ATP (B) or ADP (C) for 24 h. In addition, equal amounts of ATP and ADP were incubated with the reaction buffer (without protein) for 24 h and indicated as 24 h of ATP or ADP, respectively. The obtained relative intensity for ATP is shown in blue, for ADP in green and for AMP in orange. (D,E) Phenotypic characterization of the *sbtB-R_46_A* mutant. (D) Bicarbonate-dependent photosynthetic rates per chlorophyll a (Chla) of WT and the *sbtB-R_46_A* mutant as a function of increasing HCO_3_^−^ concentrations. Cells were acclimated to either high carbon (HC) or low carbon (LC) conditions (n = 3). (E) Bicarbonate affinity represented by the K_m_ (HCO_3_^−^) values of *Synechocystis* wild type (WT) and the *sbtB-R_46_A* mutant under either high carbon (HC; black bars) or low carbon (LC; gray bars) regimes. *: p<0.05.

**Table 1:** Dissociation constants for the binding of the adenyl nucleotides ATP, ADP, AMP, and cAMP to the WT-SbtB protein (Selim et al. 2018) and its variants as measured by ITC. NT: Not tested.

| Titration/Protein | $K_d$ (μM) for SbtB-WT | $K_d$ (μM) for SbtB-K<sub>40</sub>A | $K_d$ (μM) for SbtB-R<sub>43</sub>A | $K_d$ (μM) for SbtB-R<sub>46</sub>A |
| --- | --- | --- | --- | --- |
| ATP | 47.0 | 357.1 | 212.7 | No binding |
| ADP | 33.9 | 877.1 | 471.6 | 434.7 |
| AMP | 53.5 | NT | NT | 94.34 |
| cAMP | 21.3 | 66.9 | 83.3 | 31.3 |

Next, we selected the SbtB-R_46_A variant to analyze in depth the stepwise nucleotide hydrolysis via high-performance liquid chromatography-mass spectrometry (HPLC-MS) (Fig. 1B,C, and Fig. S2). Whereas the wildtype SbtB protein converted ATP efficiently into AMP with a very small detectable amount of ADP, the apyrase activity of the SbtB-R_46_A variant was strongly impaired similar to the mutated SbtB R-loop variant (C_105_S+C_110_S), which acts as a control for weak apyrase activity (Selim et al. 2023). By contrast, ADP was turned over by both SbtB-R_46_A and the SbtB-C_105_S+C_110_S variants as efficiently as by wildtype *Sc*SbtB (Fig. 1C).

Together, the *in vitro* characteristics of the SbtB variants demonstrate the importance of the T-loop residues for both ADP and ATP binding and hydrolysis, and highlight the critical role of R_46_ in ATP binding and breakdown, and further confirm a role for the R-loop in tuning nucleotide hydrolysis.

### Impact of SbtB apyrase activity on *Synechocystis* physiology

To further reveal the impact of ATP/ADP binding and hydrolysis on *Sc*SbtB function, we chose to characterize the SbtB-R_46_A variant *in vivo* as this variant was most strongly impaired in ATP/ADP binding and hydrolysis *in vitro*. Therefore, we generated a *Synechocystis* strain in which the wildtype *sbtB* gene was replaced by a gene encoding the *sbtB-R_46_A* variant. Then, we measured the bicarbonate-dependent photosynthetic oxygen-evolution of this mutant (Fig. 1D) to determine the bicarbonate affinity (Fig. 1E). The wildtype cells exhibited low bicarbonate affinity when adapted to high ambient C_i_ conditions (5% CO_2_; indicated by HC), whereas they displayed high bicarbonate affinity when cultivated under C_i_-limiting ambient conditions (0.04% CO_2_; indicated by LC), in accord with a fully induced CCM. In our previous work, we showed that an SbtB-deficient mutant exhibited constitutive high bicarbonate affinity, even when acclimated to HC conditions. In low C_i_-grown Δ*sbtB* cells, the maximal photosynthetic rate was lower than in wildtype cells, indicating an impaired function of the Calvin-Benson-Bassham (CBB) cycle (Selim et al. 2018), in agreement with the recently discovered additional function of SbtB in regulating entire CCM and glycogen synthesis in c-di-AMP dependent manner (Selim et al. 2021; Mantovani et al. 2022). Similar to the phenotype of the Δ*sbtB* mutant, the *sbtB-R_46_A* mutant was unable to change its overall bicarbonate affinity in response to the changing C_i_ conditions. Irrespective of LC- or HC-conditions, the *sbtB-R_46_A* cells displayed an affinity for bicarbonate in between the LC- and HC-acclimated wildtype cells, respectively. However, in contrast to the Δ*sbtB* mutant, the maximal photosynthetic rate of the *sbtB-R_46_A* mutant at saturating C_i_ conditions was as high as in the wildtype (Fig. 1D), implying that SbtB-R_46_A could complement the pleiotropic effect on the CBB-cycle, in agreement with its unimpaired ability to bind c-di-AMP (Fig. S3A). Immunoblot analyses also revealed that the *sbtB-R_46_A* variant could still be partially recovered from the cytoplasmic membrane under HC conditions, indicating that its interaction with SbtA was not completely abrogated (Fig. S3). Furthermore, the *R_46_A SbtB* variant complemented the defect of the Δ*sbtB* mutant in diurnal growth (Fig. S3B), which further suggests that this variant is able to fulfill additional functions of SbtB in accord with its ability to still bind c-di-AMP. Altogether, the *sbtB-R_46_A* mutant appeared specifically impaired in its ability to change the CCM activity in response to the ambient C_i_-supply, which involves SbtA function. This indicates that the ATP-binding and/or the stepwise hydrolysis by SbtB plays a crucial role in the regulation of SbtA activity.

### SbtB acts as valve plug to prevent bicarbonate leaking out of the cells

Recently, it was reported that immediately after light to dark shifts, the uptake activity of SbtA ceases, independent of the presence of SbtB (Förster et al. 2023). This argues against a role for SbtB in rapidly switching off SbtA-mediated bicarbonate uptake. The assumption that the formation of SbtA:SbtB complex is required to allosterically inhibit bicarbonate uptake is also in conflict with the observation that a tight SbtA:SbtB-AMP complex is formed under LC conditions when AMP levels increase (Selim et al., 2018; Liu et al. 2021; Fang et al. 2021; Förster et al. 2023). Rather, under LC-conditions, cells require maximum bicarbonate uptake activity and should not inhibit the activity of the main bicarbonate transporter, SbtA. The assumption of SbtB inhibiting SbtA activity under LC conditions is also inconsistent with the phenotype of the SbtB-deficient mutant, which showed growth impairment under LC (Selim et al. 2018, 2023). Our recent discovery that the nucleotide-bound states of SbtB are not static but may be interconverted via apyrase activity provides alternative explanations for the function of SbtB.

A so far overlooked possibility is that SbtB might serve as a dynamic valve plug to close the reverse diffusion of bicarbonate through SbtA, since it belongs to the secondary-active transporters, which operate by a reversible, bidirectional mechanism (Sauer et al. 2022). The cells invest much energy to maintain a Na^+^-motif force at the cytoplasmic membrane, which is required for the Na^+^-dependent HCO_3_^-^ transporters, SbtA and BicA, to accumulate bicarbonate approximately 100-1000 times against the gradient under C_i_-limiting conditions (Hagemann et al. 2021). According to these considerations, it should be beneficial for the cells to prevent HCO_3_^-^ backward diffusion. To verify this assumption, first, we measured the intracellular ATP levels within *Synechocystis* cells after shifting cells from HC (2% CO_2_) to LC conditions. During the first 3 h after the shift, the ATP levels dropped by 10%, whereas under prolonged C_i_-limitation (24 h) the ATP levels dropped to almost 50% of the original ATP levels under HC condition (Fig. 2A), in agreement with increasing AMP levels under LC conditions (Selim et al. 2018). Next, to test our hypothesis whether SbtB could act as a safety-valve, we checked the leakage of ^14^C-bicarbonate. Therefore, cells were loaded with ^14^C-bicarbonate in the light, then cells were washed and incubated in bicarbonate-free medium in the dark for 5 min to measure leaked ^14^C in the supernatant. In fact, the leakage of HCO_3_^-^ in the Δ*sbtB* mutant was almost 50% higher than the wildtype cells (Fig. 2B).

**Fig. 2:**
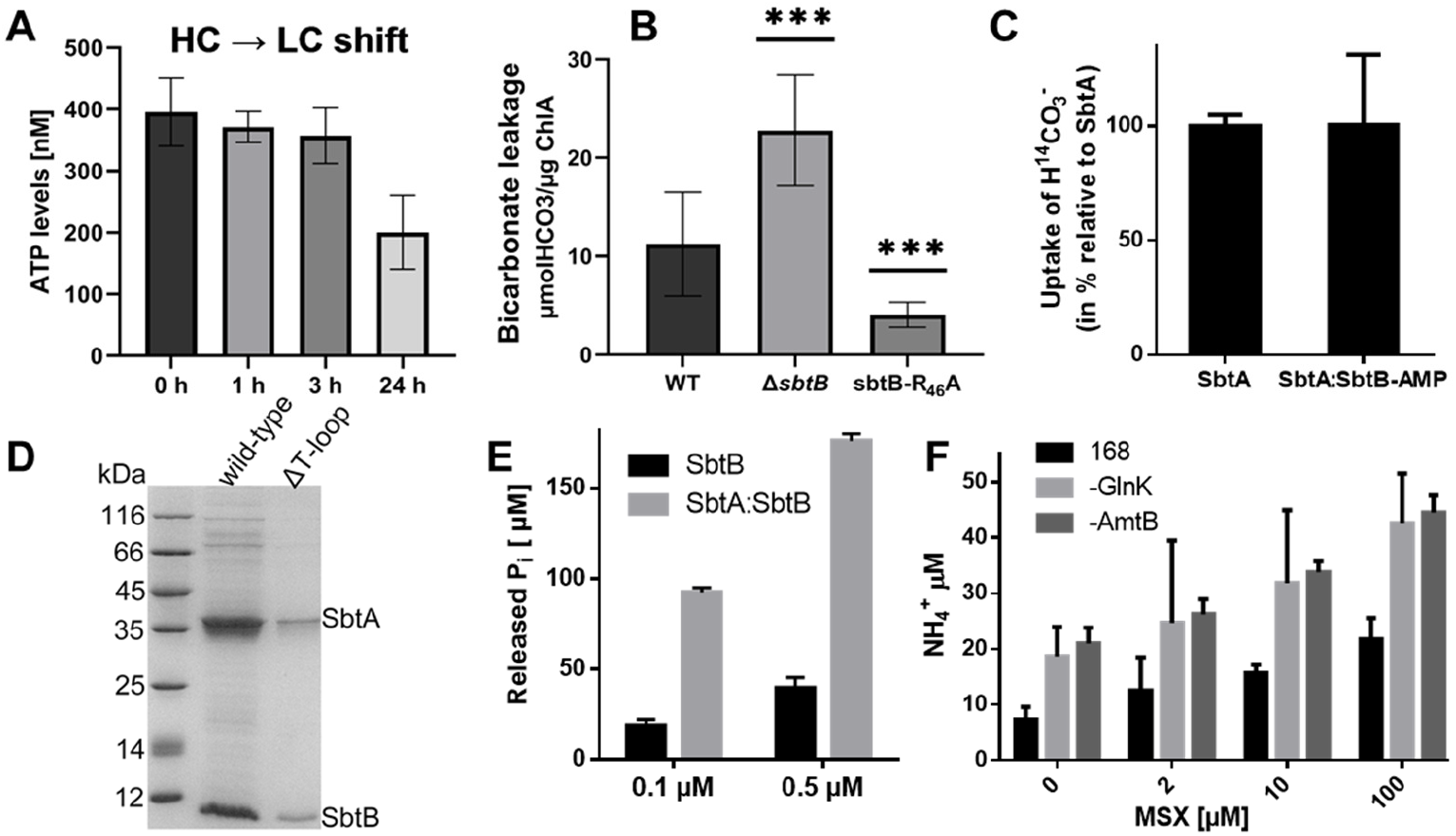
Functional characterization of PII superfamily proteins *Sc*SbtB and *Bs*GlnK. (A) Intracellular ATP levels in *Synechocystis* cells after shift from HC (0) to LC (1, 3 and 24 h) conditions. (B) Physiological investigation of SbtB function as a plug. Levels of leaked ^14^C-bicarbonate from WT, Δ*sbtB* and *sbtB-R_46_A* mutant cells. ***:p<0.001. (C) Transport activity assays of SbtA or SbtA:SbtB complex in the presence of 1 mM AMP. The assays were performed in *E. coli* membrane vesicles using ^14^C-bicarbonate. The vesicles from *E. coli* transformed by the empty pET28a plasmid were used as the negative control. (D) Interaction of SbtB ΔT-loop with SbtA compared to WT-SbtB. (E) Apyrase activity of SbtB alone or in complex with SbtA via phosphate release assay at 0.1 and 0.5 µM. (F) Level of leaked NH_4_^+^ from *B. subtilis* WT (strain 168), Δ*glnK* and Δ*amtB* mutants after treating the cells for 5 min with different concentrations of MSX, as indicated, to inhibit the GS activity to increase the intracellular levels of ammonia. For all graphs, the means and standard deviations were calculated and the data are presented as means ± SD.

All previous structural and biochemical data imply that the ATP-bound state of SbtB destabilizes the SbtA:SbtB interaction, while the AMP bound state stabilizes the interaction (Selim et al. 2018, 2023; Fang et al. 2021; Liu et al. 2021). Therefore, we assumed that the *sbtB-R_46_A* variant would be preferentially in the AMP-bound state, based on its inability to bind and hydrolyze ATP (Fig. 1B, Fig. S1, and Table 1). To test how this affects bicarbonate leakage in the *sbtB-R_46_A* mutant, we performed the ^14^C-bicarbonate leakage assay. As expected, the *sbtB-R_46_A* strain showed less HCO_3_^-^ leakage than the wild type, implying that the SbtB-R_46_A variant more efficiently prevents the backward diffusion of HCO_3_^-^ through the SbtA tunnel, consistent with the constitutive AMP bound state of SbtB-R_46_A and its inability to bind and hydrolyze ATP (Fig. 1B, and Fig. S1). These results are also in accordance with the membrane localization of SbtB-R_46_A variant (Fig. S3C). Collectively, these results generally support our hypothesis that SbtB prevents backwards HCO_3_^-^ transport. Moreover, the leakage experiments suggest that SbtB in wildtype cells dynamically changes between different states of interaction with SbtA depending on the adenyl-nucleotide bound state, which would influence the T-loop conformations to regulate the HCO_3_^-^ tunnel but not the transport activity of SbtA.

### SbtB does not influence the SbtA transport activity negatively

Recently, we revealed the mode of action of SbtA transport in presence of SbtB (PDB: 7CYF), which resembles an elevator alternating-access transport mechanism. In the SbtA:SbtB complex, when the SbtB T-loop is inserted into the inter-domain cleft of SbtA, it partially blocks the cytoplasmic substrate exit tunnel of SbtA and induces the cytoplasmic inward open conformation of SbtA by causing the SbtA subunits (core and gate domains) to undergo a significant rigid-body movement against each other (Liu et al. 2021). Intriguingly, the SbtB T-loop residue R_46_ has a dual function in stabilizing both SbtB-ATP and SbtA:SbtB interactions (Fang et al. 2021; Selim et al. 2023; Fig. S1). Based on the ability of the *sbtB-R_46_A* variant to localize to the membrane (Fig. S3C), we assumed that SbtB would still interact partially with SbtA even when the SbtB T-loop cannot be inserted firmly in SbtA. This complex would resemble the SbtA:SbtB complex with SbtB in the ATP bound state, and would correspond to either an SbtA-occluded conformation, in which the substrate tunnel is inaccessible from the intracellular or extracellular space, or to SbtA in fully open conformation, allowing HCO_3_^-^ leakage. By contrast, when the SbtB T-loop is deeply inserted in SbtA and the SbtA:SbtB interaction is stabilized via AMP or ADP, the SbtB-AMP or SbtB-ADP bound states would not influence the inward transport activity but rather prevent the back-flux of accumulated HCO_3_^-^. Those assumptions were further validated experimentally by performing *in vitro* transport activity assays in membrane vesicles containing either SbtA alone or SbtA:SbtB complex in presence of AMP to promote the SbtA-inward state (Liu et al. 2021). The activity assays showed that the SbtB-AMP state in the complex does not influence on the HCO_3_^-^ transport activity (Fig. 2C), compared to that of free SbtA, supporting our conclusions that SbtB doesn’t block bicarbonate import by SbtA but only prevents the backward HCO_3_^-^ flux (Fig. 2B). Also, we tested a T-loop variant (SbtB ΔT-loop), in which we deleted the entire T-loop so that the T-loop cannot be inserted into SbtA to determine whether the SbtB ΔT-loop variant can still interact with SbtA, resembling either the SbtA-occluded or the HCO_3_^-^ leakage state(s). Indeed, the SbtB ΔT-loop variant was still able to complex with SbtA, despite its weak interaction compared to wildtype *Sc*SbtB (Fig. 2D). This further explains a previous result showing that in absence of AMP, SbtB was able to complex with SbtA in the inward active state, where the HCO_3_^-^ substrate was clearly defined in SbtA when the SbtB T-loop was disordered (PDB: 7EGL) (Fang et al. 2021).

### Structural bases for the role of SbtB in preventing the leakage

In order to get deeper insights into the T-loop independent SbtB:SbtA complex, which likely could resemble the weak SbtA:SbtB-ATP complex, we solved the structure of SbtA:SbtB ΔT-loop complex at 3.1 Å resolution by cryo-EM (Fig. S4). In the SbtA:SbtB ΔT-loop structure SbtA, forms a membrane-localized homotrimer with symmetric orientation towards the homotrimeric SbtB ΔT-loop facing from the cytoplasmic side (Fig. 3A). Without the T-loop, SbtB can still interact primarily with the cytoplasmic phase of SbtA via van der Waals’ interactions to form a heterohexameric complex of trimer-trimer interphase, but the interactions are restricted mainly to TM2 and TM9 of SbtA and two loops of SbtB. Specifically, a salt bridge between Arg_333_ on TM9 and Glu_13_ on the loop between α1 and β1 of SbtB, as well as a hydrogen bond between Arg_333_ and Tyr_87_ on the loop between α2 and β4 (B-loop) could be found in the complex. In addition, Lys_14_ between α1 and β1 of SbtB could also form a salt bridge with Glu_40_ on TM2 of SbtA (Fig. 3A). It is noteworthy that this salt bridge is formed due to the more condensed architecture of SbtA:SbtB ΔT-loop complex, which cannot be found in our previously reported SbtA:SbtB-AMP complex (PDB: 7CYF), where the T-loop is ordered and inserted into SbtA.

**Fig. 3:**
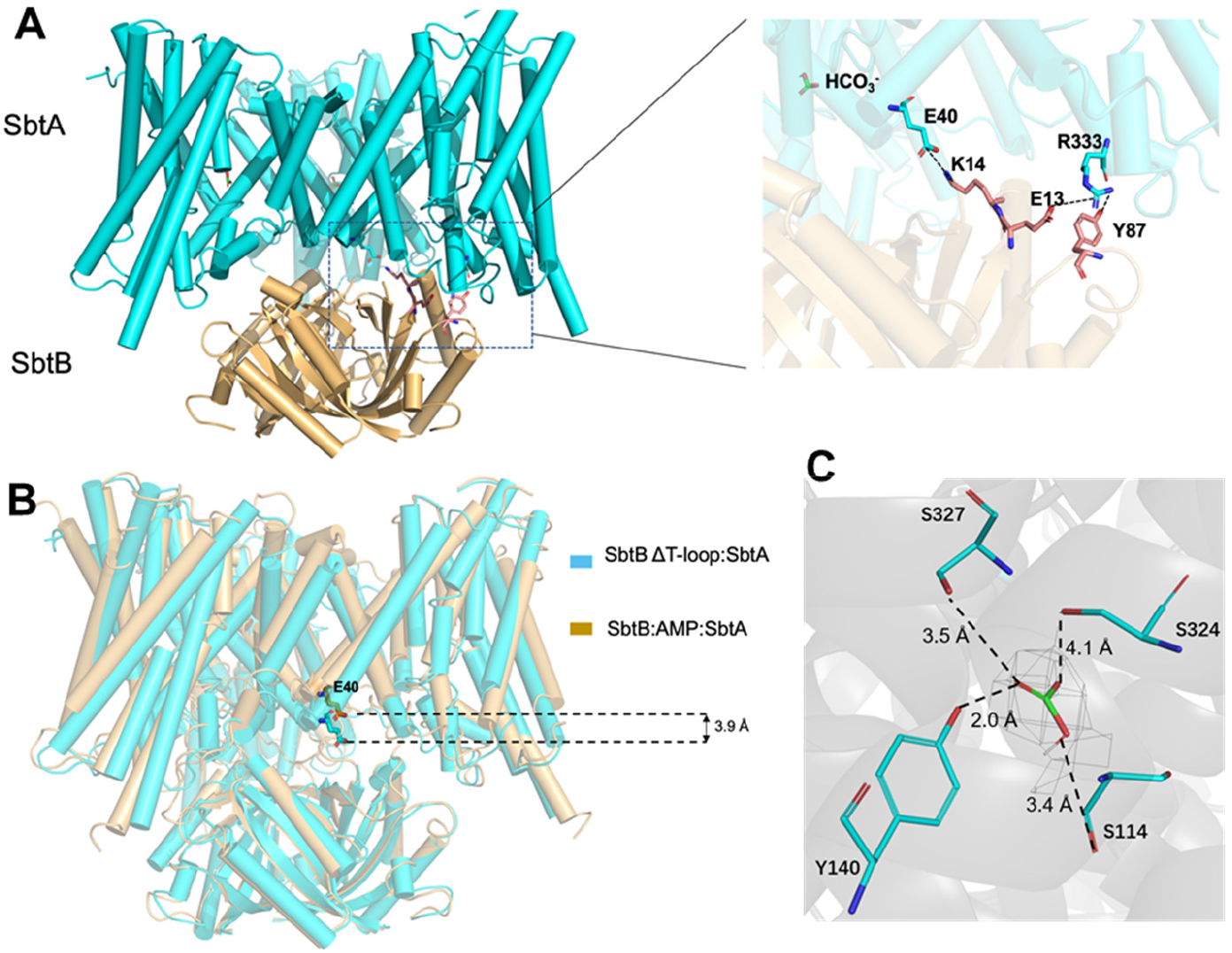
cryo-EM structure of SbtA:SbtB ΔT-loop complex. (A) Overall structure of SbtA:SbtB ΔT-loop complex. SbtA is colored cyan and SbtB is colored brown. Inset in (A): A zoom-in displaying the interface of SbtA and SbtB. Interacting residues are shown as sticks. The salt bridge and hydrogen bonds are indicated as dotted line. (B) Superposition of SbtA:SbtB ΔT-loop complex and SbtA:SbtB-AMP complex (PDB:7CYF). SbtB ΔT-loop approaches to SbtA for ∼3.9 Å measuring E_40_. (C) The HCO_3_^−^ binding site. The HCO_3_^−^ molecule is shown as sticks and colored by atoms. The polar interactions are indicated by dashed lines. The cryo-EM density map of HCO_3_^−^ is shown in gray mesh at 5 σ.

In fact, superposition of the two complexes yield an RMSD of 2.19 Å. While SbtB could almost be perfectly superposed to the previously obtained structure, SbtB ΔT-loop is approaching to SbtA by only about 3.9 Å (measuring Glu_40_), which seems more compact and reminds of the elevator mechanism (Fig. 3B), probably due to the absence of the T-loop. This approach of SbtB ΔT-loop toward SbtA led to the relative shift between the gate domain and the core domains of SbtA. This shift causes the core domain of SbtA within the SbtA:SbtB ΔT-loop complex to be uplifted, while the gate domain in this complex (TM6b and the loop between TM6b-TM7) was immobile and superimposed perfectly with our previous SbtA:SbtB-AMP complex (PDB: 7CYF) (Fig. 4A). As a result, the HCO_3_^-^ and Na^+^ binding sites are relatively elevated by about 4.1 Å (measuring Ser_114_) and 3.6 Å (measuring Phe_110_), respectively. Surprisingly, near the SbtA Ser_114_, which was reported to form the HCO_3_^-^ binding site (Liu et al. 2021; Fang et al. 2021), a clear electron density that perfectly matches HCO_3_^-^ was observed (Fig. 3C, and Fig. S4E), despite the fact that no HCO_3_^-^ was added during the purification. Although our buffer contained 300 mM Na^+^, the Na^+^-binding site did not appear occupied, supporting the notion that SbtA indeed possesses higher affinity towards HCO_3_^-^ than to Na^+^. The bicarbonate binding cavity is located at the interface between the gate domain and the core domain and interacts with Ser_114_, Tyr_140_, Ser_324_, Ser_327_ (Fig. 3C).

**Fig. 4:**
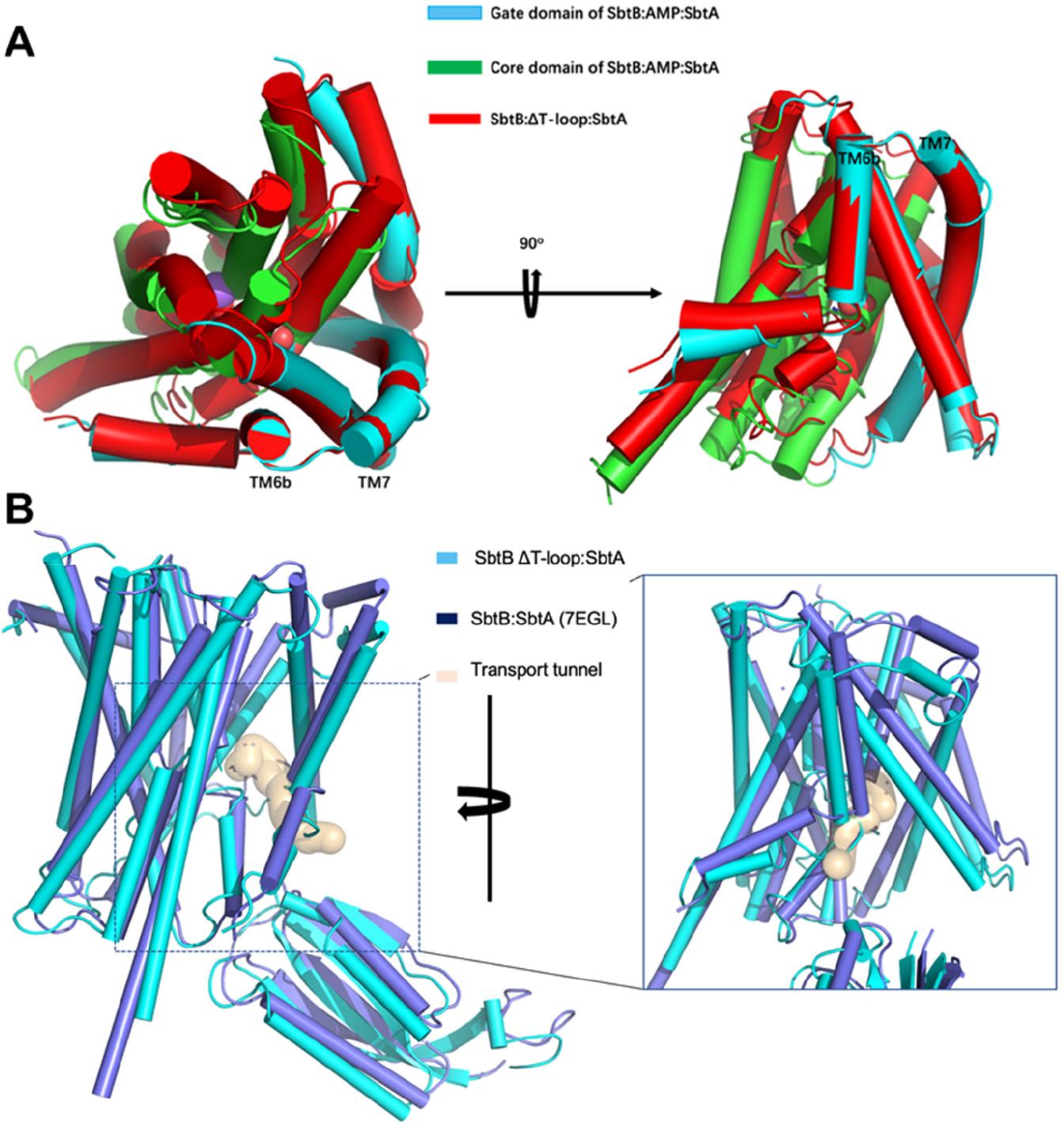
Structural comparison between current and previous SbtA:SbtB complexes. (A) Superposition of SbtA from SbtA:SbtB ΔT-loop complex and SbtA:SbtB-AMP complex (PDB:7CYF), respectively. The core domain and gate domain of SbtA (7CYF) are colored differently as indicated. (B) A simulated tunnel starting from the binding site of HCO3-in SbtB:SbtA complex (PDB:7EGL). The superposition revealed that this tunnel is blocked by the loop between TM6a and TM6b of SbtA in the SbtA:SbtB ΔT-loop complex. The tunnel is simulated by Mole 2.

We recently showed that the structure of free SbtA represents an occluded conformation (Liu et al. 2021). The superimposition of SbtA:SbtB ΔT-loop complex over the free SbtA (PDB: 7CYE) showed little difference and yielded an RMSD of 0.8 Å, indicating that the SbtA:SbtB ΔT-loop complex is also in the occluded conformation, where HCO_3_^-^ binding site is inaccessible from either intracellular or extracellular side and the HCO_3_^-^ molecule is trapped inside the SbtA tunnel. Moreover, the superimposition of SbtA:SbtB ΔT-loop complex over the SbtA:SbtB-AMP (PDB: 7CYF) complex, where the HCO_3_^-^ binding site is accessible from the intracellular space, revealed that the gate and core domains of SbtA in the SbtA:SbtB-AMP complex undergo a significant rigid-body movement against each other. As a result, the core and gate domains of SbtA in the SbtA:SbtB-AMP complex (PDB: 7CYF) separate from each other keeping the complex in an inward-open conformation toward the cytoplasm (Fig. 4A). By contrast, the SbtA:SbtB ΔT-loop complex returns back to the original occluded state exhibited by free SbtA (PDB: 7CYE) (Fig. 4). Therefore, the SbtA:SbtB ΔT-loop complex structure could represent a snapshot of the leaked HCO_3_^-^ trapped inside the SbtA tunnel when the SbtB T-loop is disordered. In this structure representing the occluded SbtA conformation of the HCO_3_^-^ transport cycle, the intermolecular interactions within the complex are restricted to the SbtA core domain due to the absence of the SbtB T-loop (Fig. 3A). Of note, a close examination of the SbtA:SbtB complex (PDB: 7EGL) solved in absence of AMP by X-ray crystallography, where the SbtB T-loop was also disordered, revealed SbtA in an inward facing state and the HCO_3_^-^ binding site in an accessible state from the intracellular space. Moreover, we simulated a tunnel starting from the binding site of HCO_3_^-^ in this complex (PDB: 7EGL), and the tunnel is clearly blocked by the loop between TM6a and TM6b of SbtA in SbtA:SbtB ΔT-loop complex (Fig. 4B). This strongly supports that SbtB undergoes dynamic structural changes in the complex to regulate the SbtA transport activity. This interpretation solves the paradox of SbtA:SbtB complex formation under low C_i_-conditions: instead of inhibiting bicarbonate uptake, SbtB prevents its leakage, as in the SbtA:SbtB-AMP complex, the cytoplasmic HCO_3_^-^ tunnel entry of SbtA is partially closed (PDB: 7CYF).

### The presence of oxidized SbtB in the SbtA:SbtB complex further supports SbtB’s apyrase activity

In light of the above results and their implications, the redox-regulated nucleotide hydrolysis activity of SbtB makes perfect sense, as it induces the tight SbtA:SbtB-AMP complex in the oxidized (dark) state. Within all reported SbtA:SbtB complexes (PDBs: 7CYF, 7EGL and 7EGK), the R-loop of SbtB forms the disulfide bridge including our recent SbtA:SbtB ΔT-loop complex (PDB: 7X1Q). We used the phosphate release assay to test the possibility that the apyrase activity of the oxidized SbtB might be further modulated by the interaction with SbtA. Indeed, we observed ≈ 4 fold increased phosphate release for SbtB in complex with SbtA as compared to SbtB alone (Fig. 2E). Finally, we assessed whether the reduced form of SbtB would still interact with SbtA. The experiments demonstrated that the reduced state mimicking variants (SbtB-Δ104 and SbtB-C_110_S) could still interact with SbtA in presence of AMP and dissociate in presence of ATP similar to wildtype SbtB (Fig. S5).

These results imply that the SbtB-redox switch does not influence the interaction with SbtA but rather accelerates the hydrolytic activity of SbtB in response to the cellular redox-state. Altogether, the formation of weak complexes of SbtB with SbtA in presence of ATP or of the SbtB ΔT-loop variant (Fig. 2) suggests that a fraction of SbtB is always close to SbtA in a sort of stand-by to open/close the SbtA tunnel.

### GlnK is a valve plug for AmtB in *Bacillus subtilis*

Intriguingly, a paradox similar to what is described here for SbtA:SbtB interaction was previously reported for the canonical PII protein GlnK in *Bacillus subtilis*. In *E. coli*, GlnK was shown to interact and inhibit the ammonium transporter AmtB (Conroy et al. 2007). In *B. subtilis, Bs*GlnK is highly expressed when grown with a poor nitrogen source, for example, with nitrate as sole nitrogen source, which is slowly reduced to ammonia for its assimilation by glutamine synthetase (GS) (Heinrich et al. 2006; Forchhammer et al. 2022; Forchhammer & Selim 2020). Localization experiments revealed that *Bs*GlnK is AmtB-associated under poor nitrogen supply (Detsch & Stülke 2003; Kayumov et al. 2011), when the AmtB transporter should support maximum ammonium transport. When nitrate ions are reduced to ammonia/ammonium (which are in a pH dependent equilibrium), the membrane permeable ammonia molecules may diffuse out of the cells before being assimilated by GS. Hence under nitrogen-limiting conditions, a highly active ammonium transport is required to recuse ammonia/ammonium molecules that leaked out of the cells, and therefore, GlnK and AmtB are highly induced and interact (Heinrich et al. 2006). In light of the above results, instead of inhibiting ammonia uptake, *Bs*GlnK could also act as a valve plug to prevent the leakage of ammonium through the AmtB ammonia pore, similar to the action of SbtB on SbtA.

To proof this hypothesis, we first showed that *Bs*GlnK indeed localizes to the membrane in an AmtB-dependent manner (Fig. S6) by fusing YFP to the C-terminus of *Bs*GlnK and analyzing its localization under nitrate growing conditions. In an AmtB-deficient mutant (Δ*amtB*), the *Bs*GlnK-YFP was solely cytoplasmic localized and never found membrane-associated, while in *amtB+* strain the *Bs*GlnK-YFP was found to be membrane localized, which indicates that AmtB is required for recruiting *Bs*GlnK to the membrane. Next, ammonium leakage experiments were performed with *B. subtilis* strains deficient in either *glnK* or *amtB*. To further trigger intracellular ammonium formation in nitrate-growing cells, samples were treated with increasing concentrations of the GS inhibitor methionine sulfoximine (MSX) to prevent ammonium assimilation by GS. GlnK- and AmtB-deficient cells displayed significantly higher extracellular ammonium levels than wildtype cells, clearly indicating ammonia leakage (Fig. 2F). The increased extracellular ammonium levels in AmtB-deficient cells illustrates the contribution of AmtB in counteracting ammonium leakage. Since in GlnK-deficient cells the same amount of extracellular ammonium was found as in the AmtB mutant, we can safely assume that in the absence of GlnK, AmtB cannot contribute to efficient ammonium scavenging. Apparently, in the absence of GlnK, the net uptake of AmtB is zero, which is the case if uptake equals the outward diffusion.

## Discussion

Recently, we have shown that in addition to linear-adenine nucleotides, SbtB also binds the cyclic nucleotides cAMP and c-di-AMP, and that by sensing cAMP, SbtB modulates C_i_ uptake and metabolism (Selim et al. 2018, 2021). During the day, the cells incur high-energy costs to intracellularly accumulate elevated concentrations of bicarbonate under C_i_-limiting conditions and to maintain the Na^+^-homeostasis required for HCO_3_^-^ transport to support efficient CO_2_ assimilation and to synthesize glycogen as an energy reservoir for the night phases. The present work has established another important role for SbtB especially during dark periods or prolonged C_i_-limitation. Our results clearly support the necessity of SbtB for the maintenance of high intracellular bicarbonate levels through its interaction with SbtA by preventing the outflux of the energetically expensive HCO_3_^-^. This also reduces energetically unfavorable Na^+^ influx, which is counteracted by secondary active sodium-proton antiport activity, under conditions when no further bicarbonate is needed. Hence, SbtB is able to simultaneously play multiple roles, reminiscent of the multitasking PII signaling proteins (Forchhammer et al. 2022; Forchhammer & Selim 2020; Mantovani et al. 2023). Our results further indicate that the apyrase activity of *Sc*SbtB stabilizes AMP-bound SbtB at the membrane (Selim et al. 2023), in a process whereby the R-loop of SbtB senses the differential redox-state between periods of dark and light. This prevents backward diffusion of bicarbonate (and Na^+^) out of the cells especially during night.

From the essence of the now available data on SbtB and its interaction partners, we propose the following refined model for the control of bicarbonate uptake through the SbtA:SbtB complex. In the light, when the R-loop is in a reduced state, there is competitive binding of adenine nucleotides to SbtB. During active photosynthesis, ATP is by far more abundant than AMP and ADP, whereas the concentration of cAMP depends on the CO_2_ supply to the cells rather than on the uptake of HCO_3_^-^ by CCM (Hammer et al. 2006). *In vitro*, cAMP, known as a high-C_i_ signal, has the highest affinity for *Sc*SbtB and the cAMP-bound state prevents binding of AMP, which is known as low-C_i_ signal (Selim et al. 2018). This adenyl-nucleotide competition is able to activate/inactivate the SbtA transport activity according to the C_i_ availability. At high C_i_, cyanobacteria accumulate high ATP and low AMP, while ATP levels drop and AMP levels increase under prolonged C_i_-limitation (Fig. 2A; and Selim et al. 2018). Therefore, it seems plausible that during the day, most of SbtB is either in the ATP-bound state or in the soluble cAMP-bound state at high CO_2_, while under prolonged C_i_-limitation SbtB accumulates in the ADP/AMP-bound state and localizes to the membrane (Selim et al. 2018). Under those conditions of C_i_-limitation and with help of SbtB, SbtA adapts an elevator alternating-access transport mechanism, as follows:

First, in the outward-open conformation, SbtA takes up the substrates HCO_3_^-^ and Na^+^. Binding of substrate seems to induce an SbtA-occluded conformation, as seen in our previous free SbtA structure (PDB: 7YCE), with bound Na^+^-ion. Subsequently, in the presence of AMP-bound SbtB, the SbtB T-loop inserts deeply into the inter-domain cleft of SbtA forming the SbtA:SbtB-AMP complex (PDB: 7CYF). In this complex, upon SbtB binding, the SbtA core domains move toward the cytoplasmic space against the immobile SbtA gate domains, causing the SbtA-substrate tunnel to be in an inward-open conformation toward the cytoplasm, and accordingly facilitating the release of substrates into the cytoplasm. At the same time, the SbtB T-loop partially blocks the SbtA-substrate tunnel exit, preventing the backward transport activity but not the inward transport, in agreement with our transport assays of SbtA:SbtB-AMP complex and the ^14^C-flux analysis of *ΔsbtB* mutant under C_i_-limitation (Fig. 2B,C). When low C_i_-acclimated cells are exposed to darkness, the ATP levels further drop and the SbtB apyrase activity, which seems to be even accelerated in the SbtA:SbtB complex (Fig. 2E), is further stimulated by the oxidation of the R-loop. This results in an increased population of ADP- and AMP-bound SbtB states, preventing the leakage of the HCO_3_^-^ accumulated during the day through the cytoplasmic substrate tunnel entry. The subsequent rise in ATP levels and ATP binding to SbtB reorients the SbtB T-loop in another conformation incompatible with SbtA, causing SbtA to return back to its original occluded conformation, where the HCO_3_^-^ molecule could be trapped inside the SbtA tunnel, as seen in the SbtA:SbtB ΔT-loop complex (PBD: 7X1Q). Altogether, this clearly supports our previous assumption that the transport model of SbtA is reminiscent of the elevator alternating-access transport mechanism (Liu et al. 2021), in which SbtB plays dual functions by inducing the inward conformation of the transporter and preventing the backward flux of the substrate from the cytoplasm (Fig. 5).

**Fig. 5:**
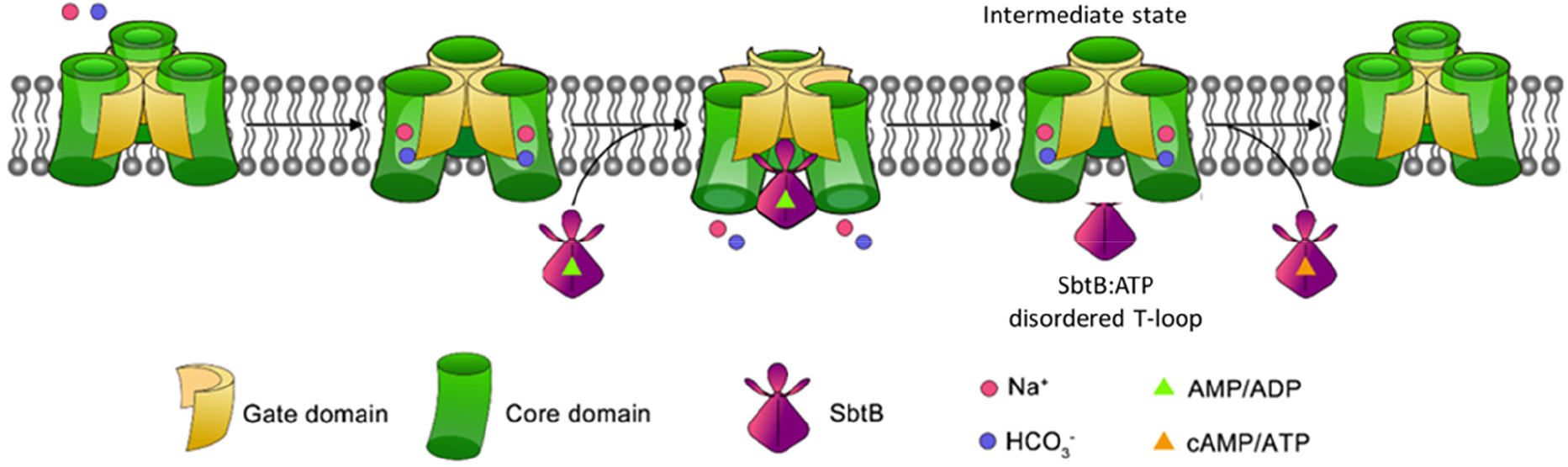
A proposed elevator mechanism of trimeric Na^+^-dependent HCO_3_^−^ transporter SbtA. SbtA in an outward-open conformation recruits the substrates Na^+^ and HCO_3_^−^ from the extracellular space This is followed by a rigid-body movement of the core domain against the immobile gate domains upon SbtB binding, a process that is stabilized by binding of AMP to SbtB, presumable also by ADP. Afterwards, the substrates are released into the cytoplasm followed by the turnover of SbtA into the resting state. SbtB can also prevent the backward flux of HCO_3_^−^ from the cytoplasm. When the T-loop of SbtB is in an incompatible conformation with SbtA, such as in ATP-bound SbtB (i.e. SbtB ΔT-loop), HCO_3_^-^ can escape from the cytoplasm as seen trapped in the tunnel of SbtA.

The structural similarity of SbtB with canonical PII proteins suggest that other proteins of the PII family might play similar roles in controlling transport tunnels. Indeed, examination of ammonium leakage by *Bacillus* clearly showed that both AmtB and GlnK are required to prevent ammonia leakage when cells are grown with poor N-source such as nitrate. The enhanced leakage in the absence of GlnK can only be reasonably explained assuming that GlnK closes the cytoplasmic face of the ammonia transporter. However, this model necessarily requires that such a complex should be dynamic like a valve plug (Fig. S7). When ammonium ion binds to the outer face of AmtB, GlnK should not rigidly block the uptake. Like in a valve-plug, the T-loop should transiently flip away from the closing position. Possibly, the previously reported ATP-ADP turnover of PII proteins (Radchenko et al. 2013) plays a role in such a valve plug activity of GlnK. The identification of a similar mechanism in the regulation of two different uptake systems in phylogenetically distant bacteria, namely, SbtA in cyanobacteria and AmtB in firmicutes, points towards a conserved function of PII superfamily proteins in the control of substrate transport. However, further research is required to solve the details of such a process.

Modelling studies revealed that reintroduction of cyanobacterial CCM, especially of the HCO_3_^-^-transporters SbtA and BicA, into plant chloroplasts could improve the carbon-assimilation rate and yield by 36-60%. However, the failure of reinstalling functional HCO_3_^-^ transporters, in particularly SbtA alone, demonstrated a lack of understanding of how those bicarbonate transporters work. In our study, we highlight the importance of considering the simultaneous reintroduction of regulatory factors such as SbtB. Remarkably, the strategy of co-expressing the *sbtAB* operon was shown to rescue the growth defect of *Corynebacterium glutamicum* carbonic anhydrase mutant under ambient air, and even more provided the wildtype cells with a growth advantage (Kirsch 2014), in contrast to *E. coli* (Du et al. 2014). In this regard, it is important to mention that the physiological meaning of SbtA:SbtB interaction in heterologous systems should be interpreted cautiously as the bioenergetics of *E. coli*, for example, is quite different from that in cyanobacteria.

Altogether, our study provides new insights into the signaling function of the PII superfamily and highlights the plasticity of PII proteins among different bacterial phyla. This is, for example, demonstrated by the ability of canonical PII to inhibit ammonium uptake and/or prevent ammonium leakage in phylogenetically distant bacteria *E. coli* and *Bacillus*, respectively.

## Material and methods

### Generation and purification of recombinant proteins

Plasmids and primers primers used in this study are listed in Supplementary (Table S1). The recombinant SbtB (wildtype or different variants [K_40_A, R_43_A, R_46_A, and C_105_S+C_110_S]) proteins from *Synechocystis* sp. PCC 6803 (*Sc*SbtB) were expressed and purified as previously described (Selim et al. 2018; Lapina et al. 2018). Truncated C-terminal *Sc*SbtB protein (SbtB-ΔC), lacking the last 6 amino acids (C_105_GPEGC_110_) at the C-terminus, was constructed as described previously (Selim et al. 2023). The *Sc*SbtB-Δ104 plasmid was used as a DNA template to generate the *Sc*SbtB-C105S+C110S and *Sc*SbtB-C110S plasmids using primer pairs contain the C_105_+C_110_ or C_110_ substitution into serine. For generation of *Sc*SbtB-K_40_A, *Sc*SbtB-R_43_A, and *Sc*SbtB-R_46_A, we used the wildtype *Sc*SbtB plasmid as a DNA template and primer pairs contain the K_40_, R_43_ or R_46_ substitution into alanine.

### Generation of Mutants

The unicellular, freshwater cyanobacterium *Synechocystis sp*. PCC 6803 was used as a reference wildtype strain in this study. The Δ*sbtB-R_46_A* mutants were generated with homolog recombination using the natural competence of *Synechocystis*, as described previously (Selim et al. 2018). For generation of Δ*sbtB-R_46_A* mutant, in which the arginine 46 (R46) is replaced by alanine (designated *sbtB-R_46_A*), a synthetic DNA fragment encoding *sbtB-R46A* with upstream and downstream regions of *slr1513* and spectinomycin resistant cassette (gBlock, IDT, USA) was cloned into BamHI-digested pUC19 vector using the Gibson assembly. The Δ*sbtB-R_46_A* mutant was selected on BG_11_ plates supplemented with spectinomycin and verified by PCR. Plasmids and primers used in this study are listed in Supplementary (Table S1).

### Protein expression and purification of SbtA-SbtB (ΔT-loop)

The codon-optimized genes *sbtA-sbtB* were synthesized by Genewiz Biotech and cloned into pET-28a (YouBio) carrying an N-terminal 6×His-tag on SbtA. The T-loop (Arg43-Ser52) truncation variant of SbtB (SbtB ΔT-loop for short) was constructed using a standard two-step PCR. The plasmids were transformed into *E. coli* C43 (WeidiBio), growing at 37°C in Luria Bertani (LB) culture medium, supplemented with 30 µg mL^−1^ kanamycin. Protein expression was induced by adding 0.4 mM isopropyl-β-D-thiogalactoside (IPTG, BioFroxx) when the OD_600_ nm reached 1.1∼1.3 for 4 h at 37°C. Next, the cells were collected and resuspended in the lysis buffer containing 25 mM Tris-HCl pH 8.0, 300 mM NaCl, 5% glycerol and stored at -80°C before use.

For purification of SbtA-SbtB (ΔT-loop) complex, the collected cells were resuspended in the lysis buffer and lysed by AH-1500 High Pressure Homogeniser (ATS, inc.) with 5 passes at 700∼800 bar. Cell debris was removed by centrifugation at 17,300 × g for 20 min. The supernatant was ultra-centrifuged at 200,000 × g for 1 h. The membrane was collected and solubilized by adding 1% (w/v) dodecyl-β-D-maltopyranoside (DDM, Bluepus) and 1% (w/v) Lauryl Maltose Neopentyl Glycol (LMNG, Anatrace) for 1 h at 4°C in the lysis buffer. After ultracentrifugation at 200,000 × g for 0.5 h, the supernatant was loaded onto a Ni-NTA resin (GE Healthcare), washed with the buffer containing 25 mM Tris-HCl pH 8.0, 300 mM NaCl, 5% glycerol, 40 mM imidazole, 0.02% (w/v) glyco-diosgenin (GDN, Anatrace) and then eluted with the same buffer supplemented with 300 mM imidazole. The eluate was then concentrated to about 1 mL by concentrators with a relative molecular mass cut-off of 10 KDa, and was applied to a Superdex 200 Increase 10/300 gel filtration column (GE Healthcare) equilibrated in the buffer containing 25 mM Tris-HCl pH 8.0, 300 mM NaCl, 5% glycerol, 0.02% (w/v) GDN. The peak fractions were collected for further procedures.

### Cryo-EM sample preparation, data collection and processing

Aliquots of 3.5 μL SbtA-SbtB (ΔT-loop) samples (∼6 mg/mL) were applied to the glow-discharged grids (Quantifoil holey carbon Cu R1.2/1.3 grid). Grids were blotted for 4.5 sec and plunge-frozen in liquid ethane vitrified by liquid nitrogen using Vitrobot Mark IV (FEI Company) at 8°C and 100% humidity.

The cryo-EM grids of SbtA-SbtB (ΔT-loop) were loaded into a Titan Krios transmission electron microscope (ThermoFisher Scientific) operating at 300 KeV with a Gatan K3 Summit direct electron detector. Totally 3,056 movie stacks were collected in a super-resolution mode with a defocus range from -1.2 to -2.0 μm. Each movie stack of 32 frames was exposed for 3.5 sec under a dose rate of 16 e/pixel/sec, resulting in a total dose of ∼55 e Å^−2^. All Krios data were collected using FEI EPU and then processed by cryoSPARC. After motion correction and contrast transfer function (CTF) estimation, a total of 1,787,952 particles were auto-picked and extracted at a 2-fold binned with a pixel size of 2.14 Å. After multi-rounds of 2D classification, 424,896 particles were used for 3D classification, searching for 3 classes using the references generated by the 3D initial model. 239,801 particles from the best class were re-extracted at a pixel size 1.07 Å without binning and used for Non-uniform 3D refinement yielding a 3.49 Å map. After multi-rounds of 3D classification, 178,612 particles were selected for final 3D refinement yielding a 3.10 Å map (Supplementary Fig. S4).

### Model building and refinement

The initial model of SbtA-SbtB complex was built by fitting the SbtA-SbtB complex (PDB: 7CYF) structures into the map using the UCSF Chimera. Then model building and refinement was accomplished manually by Coot. The final structures showed good geometry and were further evaluated using MolProbity. A list of parameters of cryo-EM data collection, processing, structure determination and refinement is provided in the Supplementary (Table S2).

### HCO_3_^-^ leakage assay

Low carbon (LC) adapted cells (WT, Δ*sbtB*, and Δ*sbtB*-R46A) were centrifuged and re-suspended in fresh BG11 medium (pH 8.0) at an OD_750_ = 1.0. Two mL of each culture were transferred to 2 mL tubes, to which H^14^CO_3_^-^ and HCO_3_^-^ were added, each with a final concentration of 200 µM. The tubes were incubated at 30 °C under a constant light with an intensity of 50 µE for 10 minutes, after which 1 mL from each sample was placed in new tubes containing 200 µL of a mix of silicone oil AR20:AR200 (Sigma-Aldrich) with a ratio of 4:1 (to separate the cells from the culture media which contains ^14C^HCO_3_^-^). The samples were then centrifuged at 12.000 rpm for 20 minutes, the supernatant was discarded and the cells re-suspended in 650 µL carbon free BG11 media (pH 8.0), after which they were incubated in darkness for 5 minutes at room temperature (to prevent use of intracellular bicarbonate and allow it to leak from the cells). 250 µL of each sample were transferred into new tubes. The tubes were then centrifuged at 12000 rpm for 1 min to remove the cells and to collect the supernatant (to determine in the supernatant the amount of released ^14^C outside the cells). In 5 mL scintillation vials (PerkinElmers), from each supernatant sample, 50 µL were added to 200 µL of 10 M formic acid (acid samples; all HCO_3_^-^ and H_2_CO_3_ are converted to CO_2_) and 50 µL were added to 200 µL of 1M NaOH (alkaline samples; all CO_2_ and HCO_3_ ^-^ are converted to H_2_CO_3_). After mixing, the samples were left to evaporate overnight at 70 °C. Finally, after evaporation, the samples were dissolved in 500 µL ddH_2_O, then 5 mL of the scintillation cocktail UltimaGold (PerkinElmers) were added to each tube, and the scintillation counts were measured in a scintillograph (Tri-Carb 2810 TR, PerkinElmers). For calculation, the acid samples were subtracted from the alkaline samples to determine the amount of radioactively labelled C_i_ that has evaporated from the solutions (which corresponds to the C_i_ that leaked from the cells). Simplified scheme is presented in (Fig. S8).

### High-performance liquid chromatography-mass spectrometry (HPLC-MS) analysis of SbtB ATPase/ADPase activity

Purified SbtB wildtype, R_46_A and C_105_S+C_110_S protein variants were dialyzed overnight at 4°C in Tris-HCl buffer [50 mM Tris/HCl, 200 mM NaCl, 1 mM MgCl_2_; pH 7.4]. 300 µM of the dialyzed enzyme was then incubated in a rotary shaker with 300 µM ATP or ADP at 28 °C in a total volume of 100 µl (as control 300 µM of either ATP or ADP was incubated in a buffer without enzyme) for 24 h. Next, the proteins were precipitated by addition of 100 µl CHCl_3_ and centrifuged for 10 min centrifugation at 21000 x g at 4 °C. Supernatants were taken and dried in a vacuum concentrator at room temperature for 2-4 h. Finally, pellets were solved in 30 µl of Millipore water and analyzed by HPLC-MS.

Nucleotide analysis was performed using an ESI-TOF mass spectrometer (micrO-TOF II, Bruker) operated in negative ion mode (mass range 85-900 m/z) and connected to an UltiMate 3000 HPLC system (Dionex). Five μl of each sample or standards (ATP, AMP or AMP) were injected into the SeQuant ZIC-pHILIC column (Merck, PEEK 150 × 2.1 mm, 5 μm) and the HPLC system was run with a flow rate of 0.2 ml/min as previously described (Kästle et al. 2015; Gratani et al. 2018) with only modified 35-minute gradient program: 5 minutes of 82% buffer A (CH_3_CN) and 18% buffer B (100 mM (NH_4_)_2_CO_3_, pH 9); 20 minutes of a linear gradient from 82% to 42% buffer A; and finally, 10 minutes of 82% buffer A. The nucleotide analysis was done using the program UmetaFlow (Kontou et al. 2023) by calculating the area under the curve (relative intensity) of the extracted ion chromatograms for ATP (505.989 m/z), ADP (426.022 m/z) and AMP (346.056 m/z). HPLC-MS experiment was performed in three biological replicates and presented in GraphPad Prism 8.4.3 as mean ±SD.

### Preparation of membrane vesicles

The right-side-out membrane vesicles were prepared according to the previous report (Liu et al. 2021). Briefly, the process was started with the preparation of osmotically sensitive *E. coli* cells. The empty plasmid or plasmids carrying SbtA or SbtA:SbtB were transformed into *E. coli* C43. Then cells were cultured in M63 medium (2 g (NH_4_)_2_SO_4_, 13.6 g KH_2_PO_4_, 0.5 mg FeSO_4_·7H_2_O, 0.246 g MgSO_4_·7H_2_O, 4.2 g KOH, 0.2% glucose, 0.1% casamino acids and 0.1 ml 0.5% vitamin B_1_ per liter) and induced with 0.4 mM IPTG at 37 °C for 4 h. The culture was centrifuged at 16000 × g to harvest the cells and the pellet was washed twice with 10 mM Tris-HCl, pH 8.0 on ice. The cells were weighted and resuspended with 30 mM Tris-HCl, pH 8.0, and 20% sucrose, and flash-frozen with liquid nitrogen.

Next, the thawed cells were diluted with 30 mM Tris-HCl, pH 8.0, and 20% sucrose (1g wet weight, per 80 mL) supplemented with 10 mM EDTA-KOH, pH 7.0 and 0.5 mg/mL lysozyme, and swirled for 30 min by means of a magnetic stirrer at room temperature. The protoplast suspensions were centrifuged at 16,000 × g for 15 min and the pellet was resuspended in 10 mL buffer containing 0.1M KH_2_PO_4_/K_2_HPO_4_, pH 6.6, 20 mM MgSO_4_ and homogenized by ULTRA-TURRAX (IKA). The suspension was poured directly into 300-fold volumes of 50 mM KH_2_PO_4_/K_2_HPO_4_, pH 6.6 supplemented with 100 μg/mL DNase and 100 μg/mL RNase. The lysate was incubated for 15 min at 37°C with vigorous swirling. Then, 10 mM EDTA-KOH was supplemented and the lysate was incubated for another 15 min. Afterwards, 15 mM MgSO_4_ was supplemented in the lysate. In the end, the lysate was centrifuged at 16,000 g for 30 min and the pellet was centrifuged at 45,000 × g for 30 min to isolate the membranes.

The isolated membranes were homogenized in a solution of 0.1 M KH_2_PO_4_/K_2_HPO_4_, pH 6.6, containing 10 mM EDTA on ice. Then, the suspension was centrifuged at 800 × g until the supernatant fluid was clear. The pellet was washed 4∼6 times by centrifugation at 45,000 × g for 30 min in the solution of 0.1 M KH_2_PO_4_/K_2_HPO_4_, pH 6.6, containing 10 mM EDTA on ice. Finally, the obtained membrane vesicles were resuspended by homogenization in the transport assay buffer^5^ (50 mM CHES-KOH pH 9.0, 0.3 mM MgSO_4_, 0.26 mM CaCl_2_, 0.22 mM K_2_HPO_4_) at a concentration of 5 mg/mL (wet weight), frozen in small aliquots in liquid nitrogen and stored in -80°C before use within a week for the best performance. The amounts of proteins in the membrane vesicles for each preparation were quantified by purifying the proteins from the same batch of cells by Coomassie brilliant blue staining.

### Transport activity assays

For each assay sample, 50 μL of membrane vesicles were thawed. The substrates of 12 mM NaCl and 0.1 nCi ^14^C labeled NaHCO_3_ (∼12 μM, American Radiolabled Chemicals) were added in the system supplemented with the transport assay buffer (50 mM CHES-KOH pH 9.0, 0.3 mM MgSO_4_, 0.26 mM CaCl_2_, 0.22 mM K_2_HPO_4_) to 100 μL. All reactions were performed at 30 4°C for 30 s and terminated by rapid filtration on a glass filter (25 mm GF/F, Whatman) by suction, followed by immediate wash of the filter with 10 mL of the washing buffer (the transport assay buffer supplemented with 120 mM NaHCO_3_ to prevent back flow). The filters were soaked in 3 mL ULTIMA Gold (PerkinElmer) overnight before liquid scintillation counting. The vesicles prepared with induced *E. coli* transformed by the empty plasmid pET-28a were tested as the control group.

### ATPase activity assays

ATPase activities were measured using the ATPase colorimetric Assay Kit (Innova Biosciences) in 96-well plates at OD_630_ nm. The protein was added in the reaction buffer consisting of 25 mM Tris–HCl pH 8.0, 50 mM KCl, 2 mM MgCl_2_, and 2 mM ATP (Sigma) to 80 μL as one reaction sample. Reactions were performed at 37°C and the amount of released phosphate group (P_i_) was quantitatively measured using a SpectraMax iD5 Multi-Mode Microplate Reader (Molecular Devices). The control groups in the absence of proteins were subtracted as background for each data point.

### Isothermal titration calorimetry (ITC)

ITC experiments were performed as previously described (Selim et al. 2018, 2019) at 20°C using a VP-ITC microcalorimeter (MicroCal) or Malvern-Microcal PEAQ-ITC in Tris-HCl buffer pH 7.9 and 150 mM NaCl buffer pH 8.0. For determination of binding isotherms, *Sc*SbtB wildtype or mutant variants (30 µM trimeric concentration) were titrated against ATP, ADP, AMP, and cAMP, as indicated.

### ATP quantification

For quantification of intracellular ATP levels, *Synechocystis* sp. PCC 6803 cells were cultivated under high CO_2_ (HC, 2% CO_2_) bubbling. A shift to low CO_2_ (LC, 0.04% CO_2_) was achieved by replacing CO_2_ with ambient air bubbling. 2 ml samples were taken at each time point and disrupted in three consecutive boiling-freezing cycles. Thereby, cells were frozen in liquid nitrogen followed by an immediate heated to 99°C under 1400 rpm agitation. After lysis, cell debris was pelleted by centrifugation at 25.000g at 4°C for 1 min. The ATP content of the supernatant was immediately measured using the ATP Determination Kit (A22066) [Molecular Probes^TM^] according to manufacturer’s instructions. Standard curves were generated using a dilution series (0 – 1000 nm) of ATP.

### NH_4_ ^+^ leakage assay

The release NH4+ into media was measured for wildtype *B. subtilis*, and both Δ*glnK* and Δ*amtB* mutants (Detsch & Stülke 2003; Kayumov et al. 2011) using either glutamate dehydrogenase (GDH) assay and phenol-hypochlorite (PH) assay. The results of both assays were comparable. In the GDH assay, GDH converts ammonia with α-ketoglutaric acid and NADPH to form L-glutamate and NADP^+^. The decrease in absorbance at 340 nm, due to the oxidation of NADPH to NADP^+^, is proportional to the ammonia concentrations in the media. The assay was performed using 200 µl of media supernatant in 100 mM phosphate buffer pH 7.4. For PH assay, a reaction mixture of 1:1:1 of solution I (10 mg sodium nitroprusside dissolve in 95.33 ml H_2_O add 4.67 ml liquefied Phenol [1.07 g/ml]), solution II (25 ml of 2.5 M NaOH, 0.7 ml NaClOH [150-180 g/liter], 74.3 ml H_2_0), and samples (media supernatant), was incubated for 15 min at room temperature and the change of the color at 625 nm was recorded and correlated to a calibration curve of standard HN ^+^ solutions.

## Acknowledgments

The project was funded by grants from the German Research Foundation (DFG) as part of the priority research program (SPP1879) to MHag (HA 2002/24-1) and to KF (Fo195/18-1), and by the Federal Ministry of Education and Research (BMBF) and the Baden-Württemberg Ministry of Science as part of the Excellence Strategy of the German Federal and State Governments to KAS (Projektförderung: PRO-SELIM-2022-14). KF, MB and KAS acknowledge the infrastructural support by the Cluster of Excellence “Controlling Microbes to Fight Infections (CMFI)” (EXC 2124–390838134). This work was supported by the Strategic Priority Research Program of the Chinese Academy of Sciences (grant numbers XDB37020301 and XDA24020302). MB gratefully acknowledge the financial support from the DFG (TRR261: project ID 398967434) and (CMFI: project ID 390838134) to Christoph Mayer. We also thank Dr. Yongxiang Gao (Cryo-EM Center) and Mr. Jishu Ren (Isotope Laboratory) at the University of Science and Technology of China for cryo-EM image acquisition and for the transport assays, respectively. The PhD thesis of PW is supported by the German Academic Scholarship Foundation (Studienstiftung des deutschen Volkes).

## Author contributions

KAS and KF conceived, initiated, and supervised the whole research; KAS, WH, MHag CZ and KF designed research; MHaf, WH, OM, RRW, KH, and MB performed research; KAS and WH analyzed data and prepared the figures; and KAS wrote the manuscript with inputs from MHag, WH, and KF. All authors approved the final version of the manuscript.

---

**Fig. S1:**
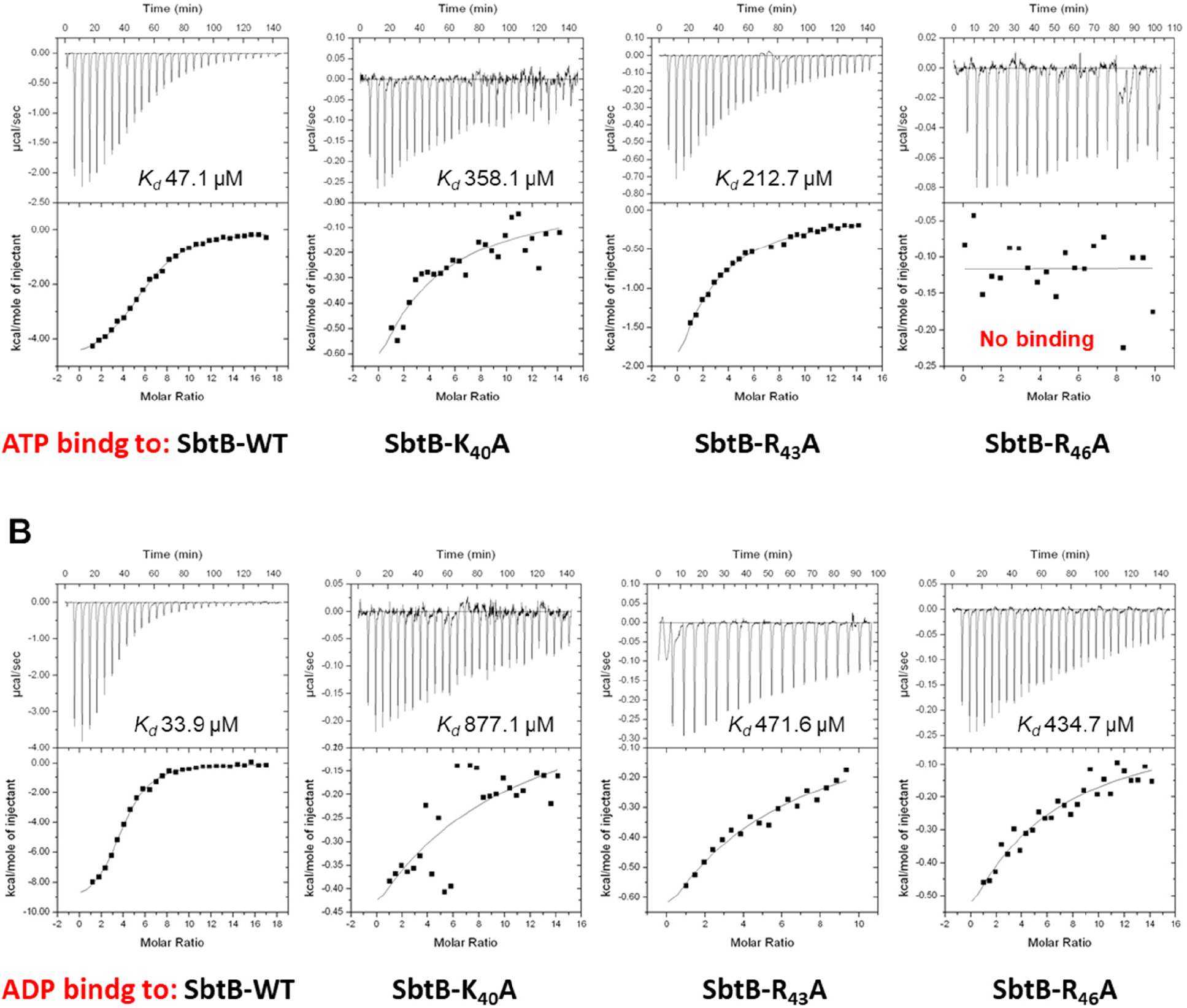
ITC analysis of SbtB and its T-loop variants (SbtB-K_40_A, SbtB-R_43_A and SbtB-R_46_A). Upper panels show the raw ITC data in the form of the heat produced during the titration of 30 µM SbtB (trimeric concentration) with ATP (A) and ADP (B); while lower panels show the binding isotherms and the best-fit curves to calculate *K_d_* values according to the one binding site models.

**Fig. S2:**
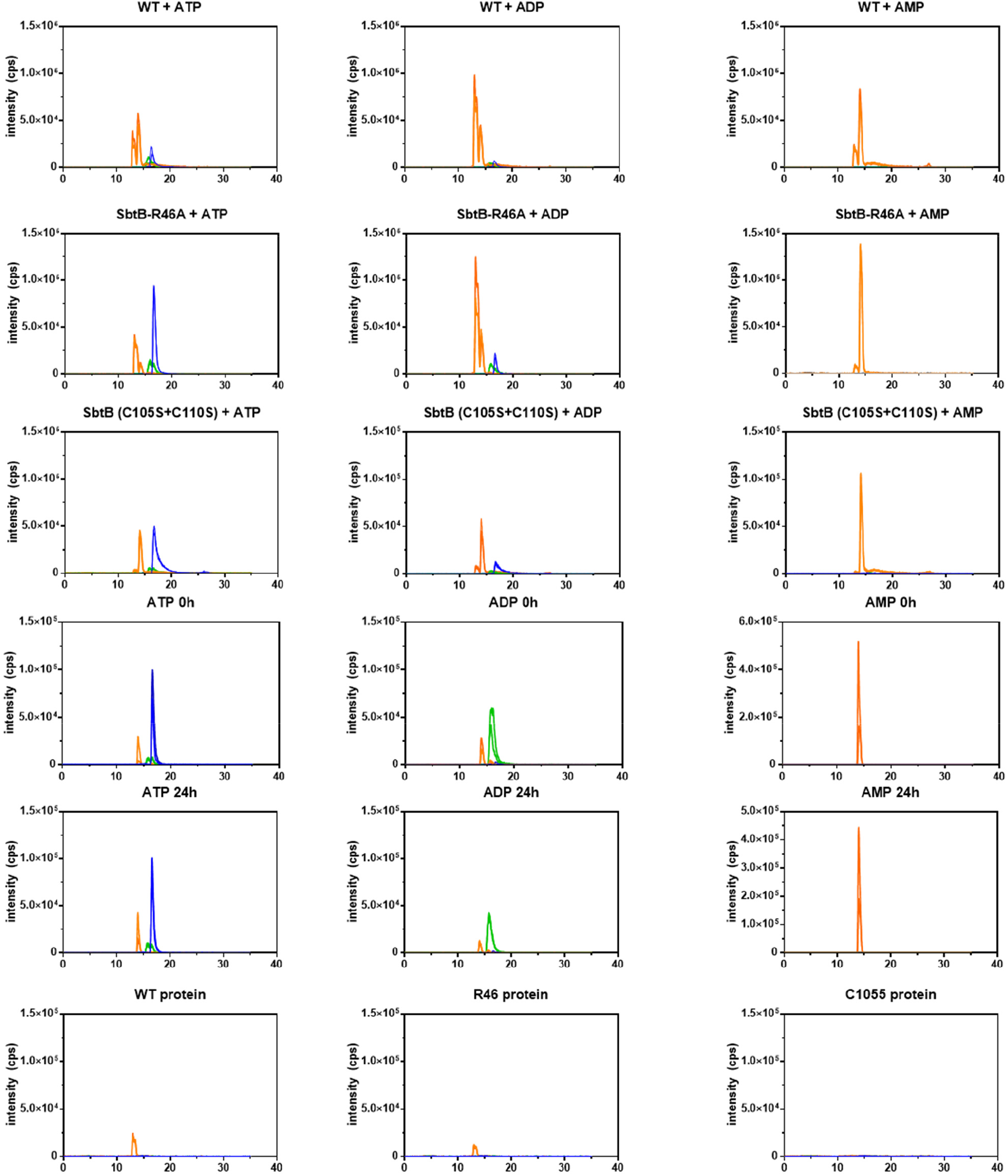
HPLC-MS analysis was performed for SbtB wildtype and its variants (R_46_A and C105S+C110S [+ve control]) and the extracted ion chromatograms for ATP, ADP and AMP were generated, as indicated. The area under the curve for each nucleotide was used to quantify relative intensity of each nucleotide.

**Fig. S3:**
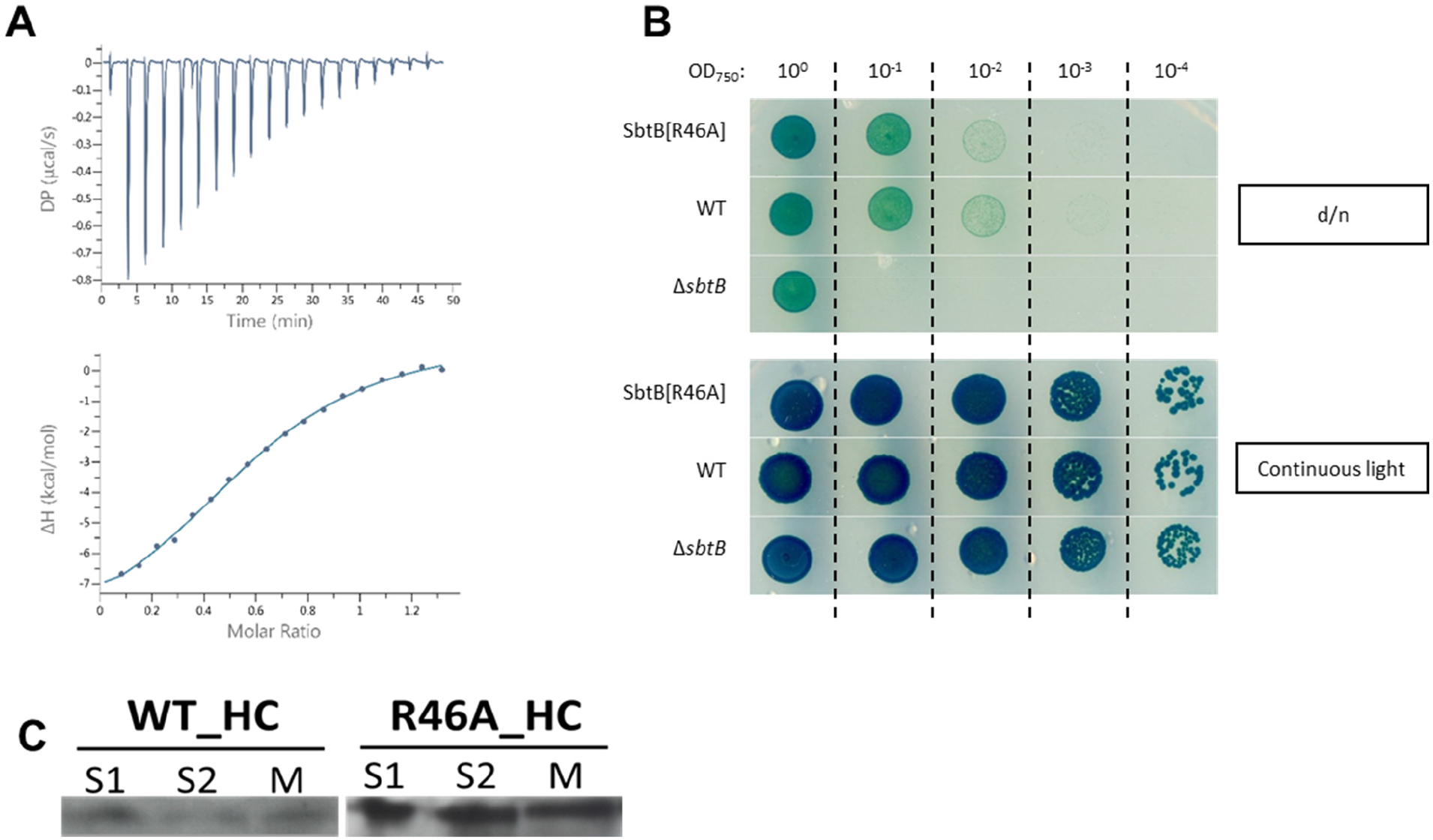
Characterization of SbtB-R_46_A mutant. (A) An isothermal calorimetric assay of recombinant SbtB-R_46_A protein (37 µM; trimeric concentration) against 750 µM c-di-AMP. (B) While Δ*sbtB* had a severe growth defect in a diurnal rhythm, the *sbtB-R_46_A* mutant did not show impaired growth compared to wildtype *Synechocystis* (upper picture). Cells were normalized to an OD_750_ of 1.0 and serial diluted to an OD_750_ of 1*10^-4^. Plates were either exposed to a diurnal rhythm (upper picture) or to continuous light (lower picture). (C) Analysis of SbtB localization. Localization of SbtB in soluble (S1 & S2) and membrane (M) fractions of cells grown under high carbon (HC; 5% CO_2_) as assessed by immunoblot using anti-SbtB antibodies.

**Fig. S4:**
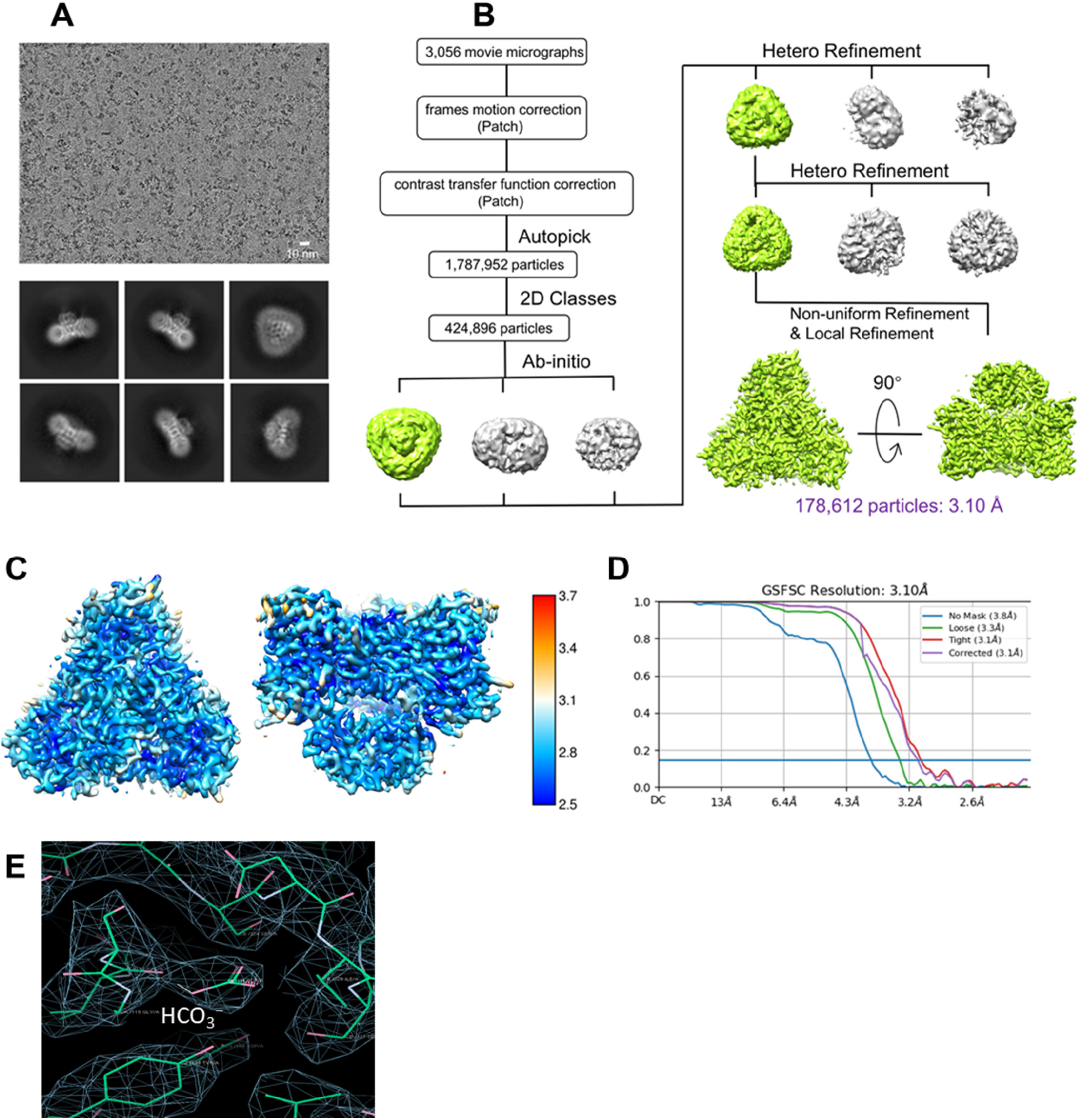
Cryo-EM analysis of SbtA:SbtB ΔT-loop complex of *Synechocystis* sp. PCC 6803. (A) Representative cryo-EM micrograph of SbtA:SbtB ΔT-loop complex and 2D classes. (B) Data processing flowchart with particle distributions. (C) Resmap resolution slice and resolution map for SbtA:SbtB ΔT-loop complex, shown in the top view and the side view espectively. (D) Fourier shell correlation (FSC) curves showing a resolution of 3.1 Å. (E) The electron density of HCO_3_^−^.

**Fig. S5:**
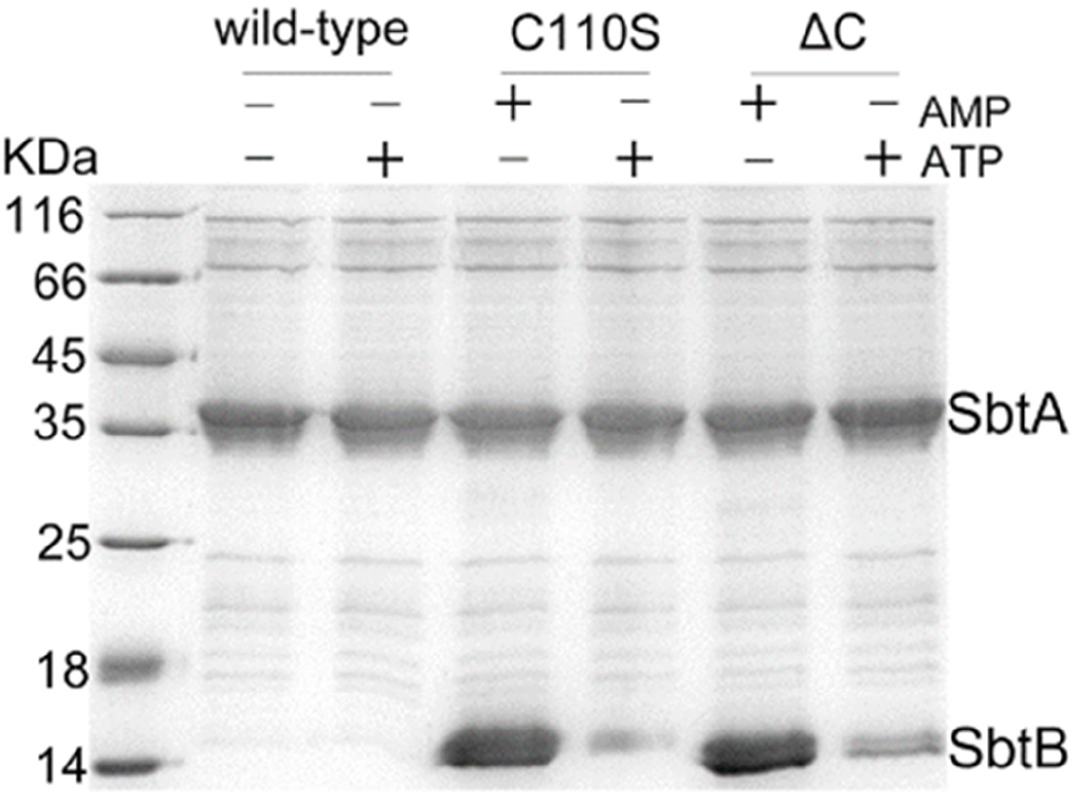
Interaction of SbtA with SbtB and its variants (C105S and ΔC, in which the entire R-loop from C_105_-C_110_ was deleted) by pulldown assays in presence and absence of ATP or AMP as indicated.

**Fig. S6:**
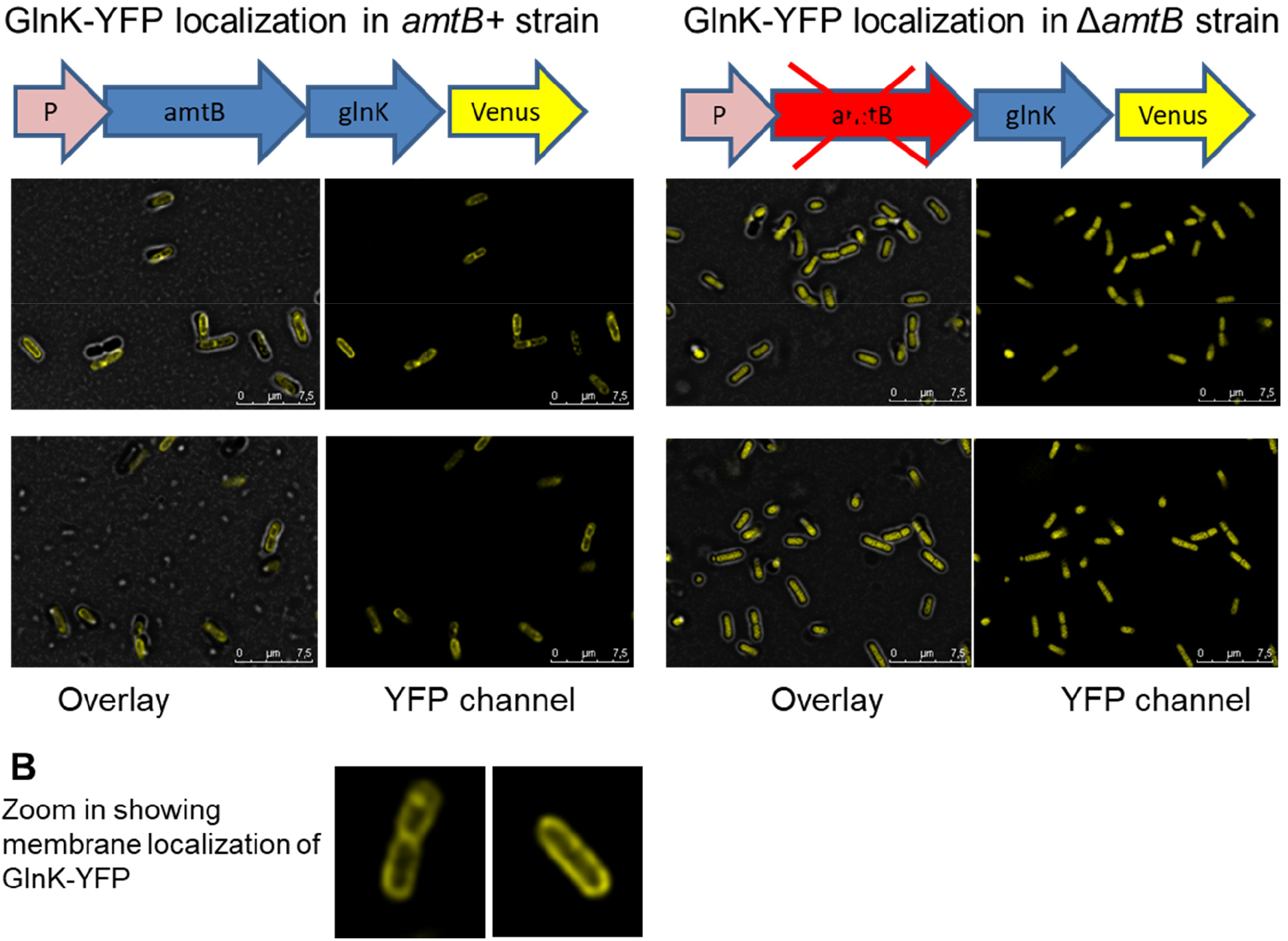
GlnK localization in *amtB+* and Δ*amtB* strains under nitrate growth conditions as indicated.

**Fig. S7:**
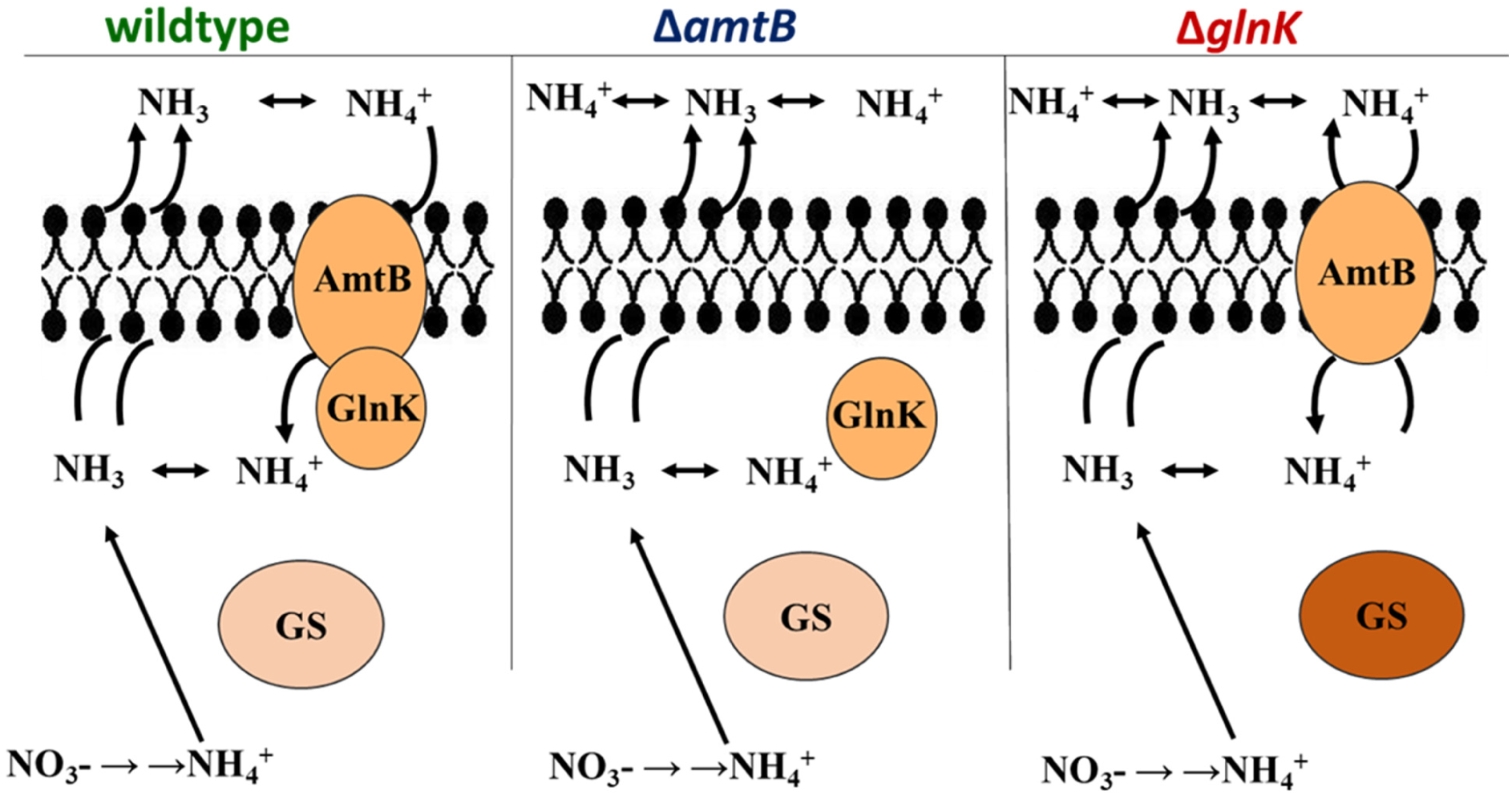
Proposed mechanism for GlnK to act as a valve plug to control the ammonium transporter AmtB under poor nitrogen conditions in *Bacillus*.

**Fig. S8:**
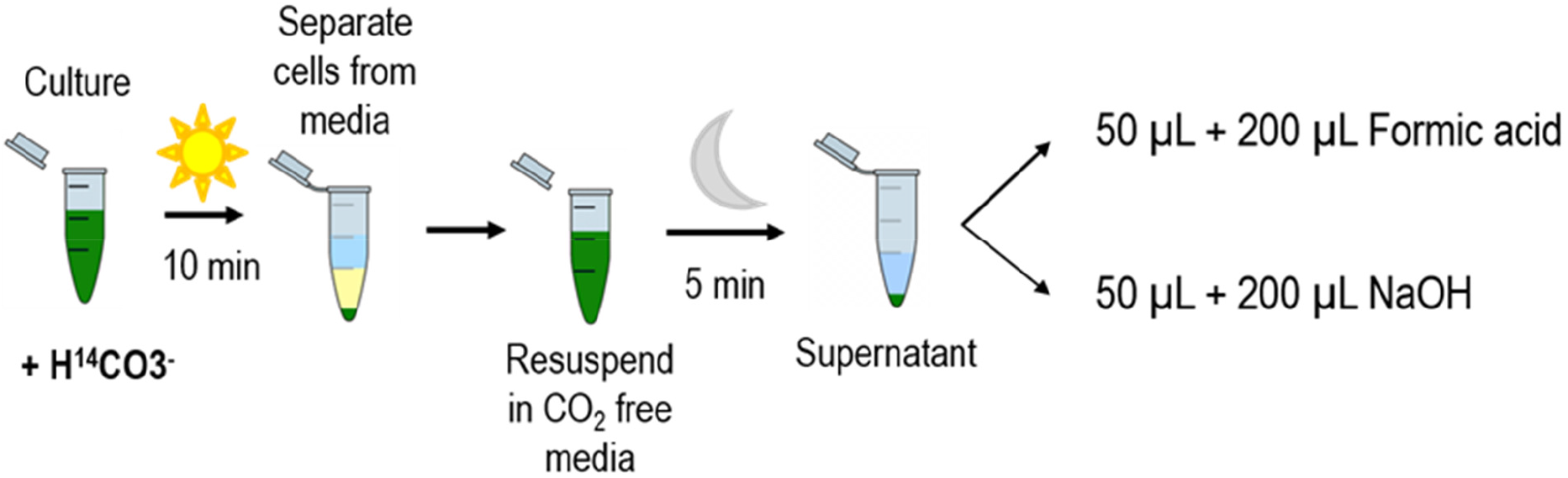
Simplified scheme of HCO_3_^-^ leak experiment protocol.

**Supplementary Table S1.**
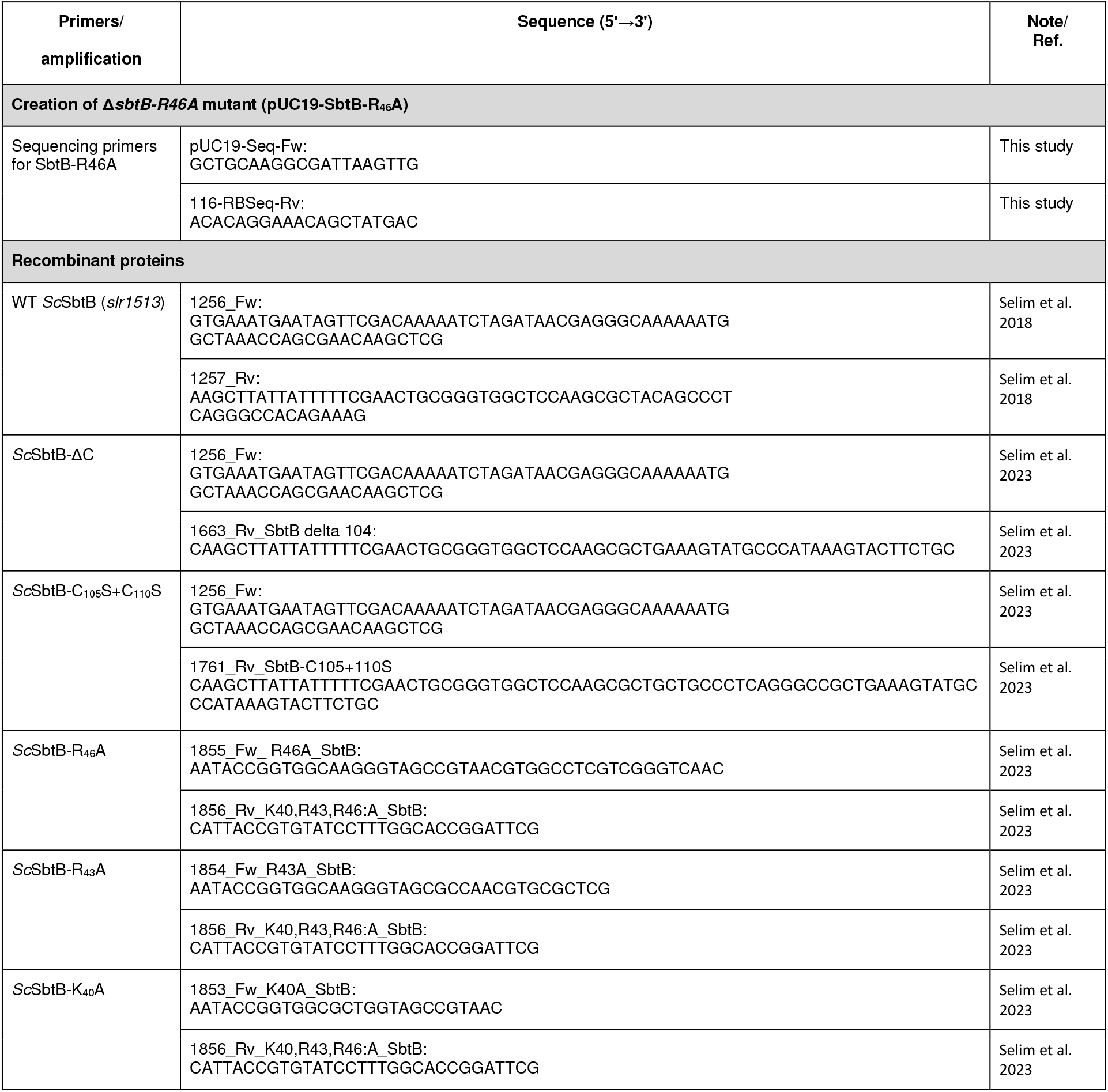
Primers and Plasmids List.

**Supplementary Table 2.**
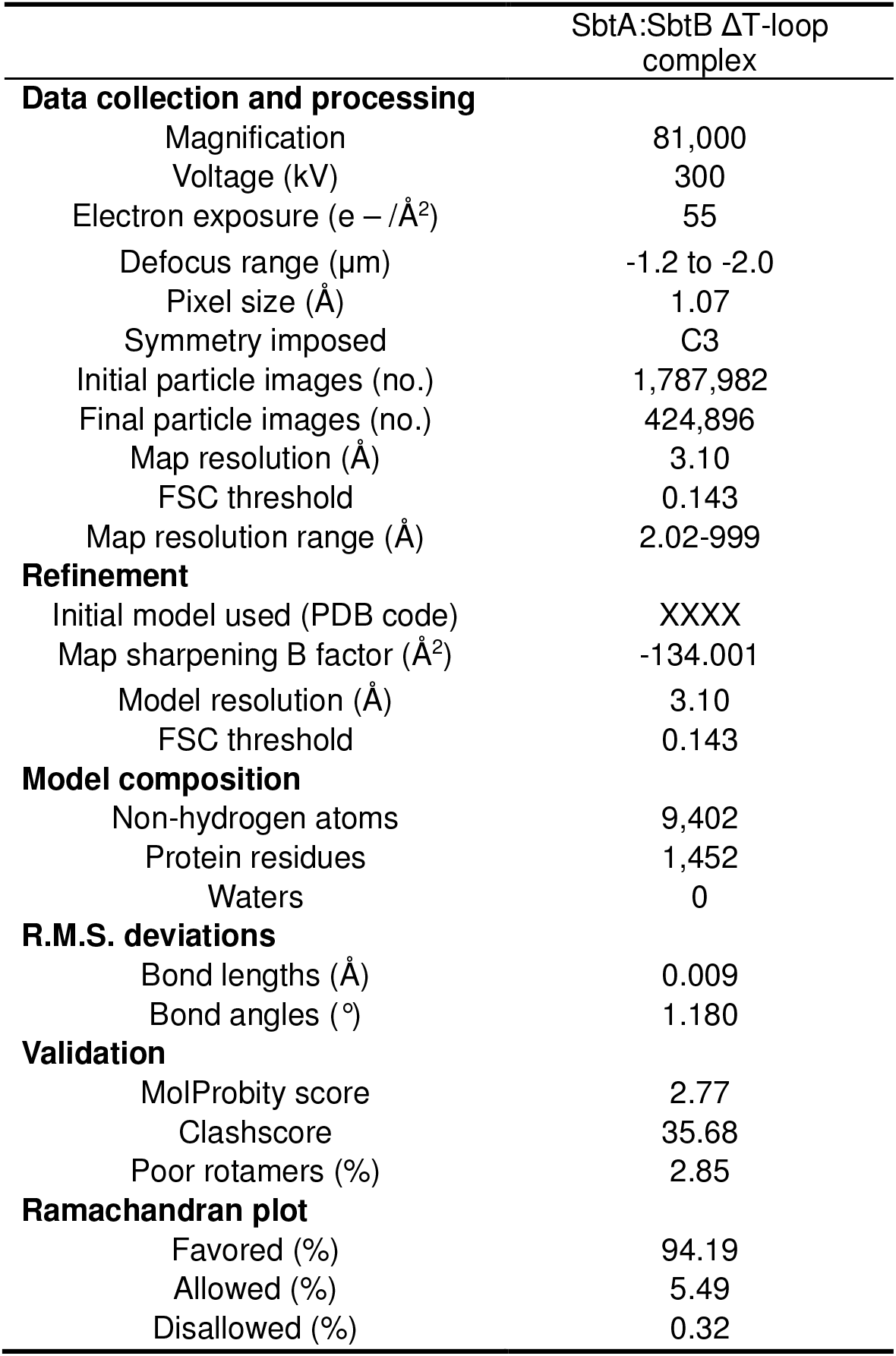
Cryo-EM parameters, data collection and refinement statistics.

