## Supplementary Information for "PII signal transduction superfamily acts as a valve plug to control bicarbonate and ammonia homeostasis among different bacterial phyla"

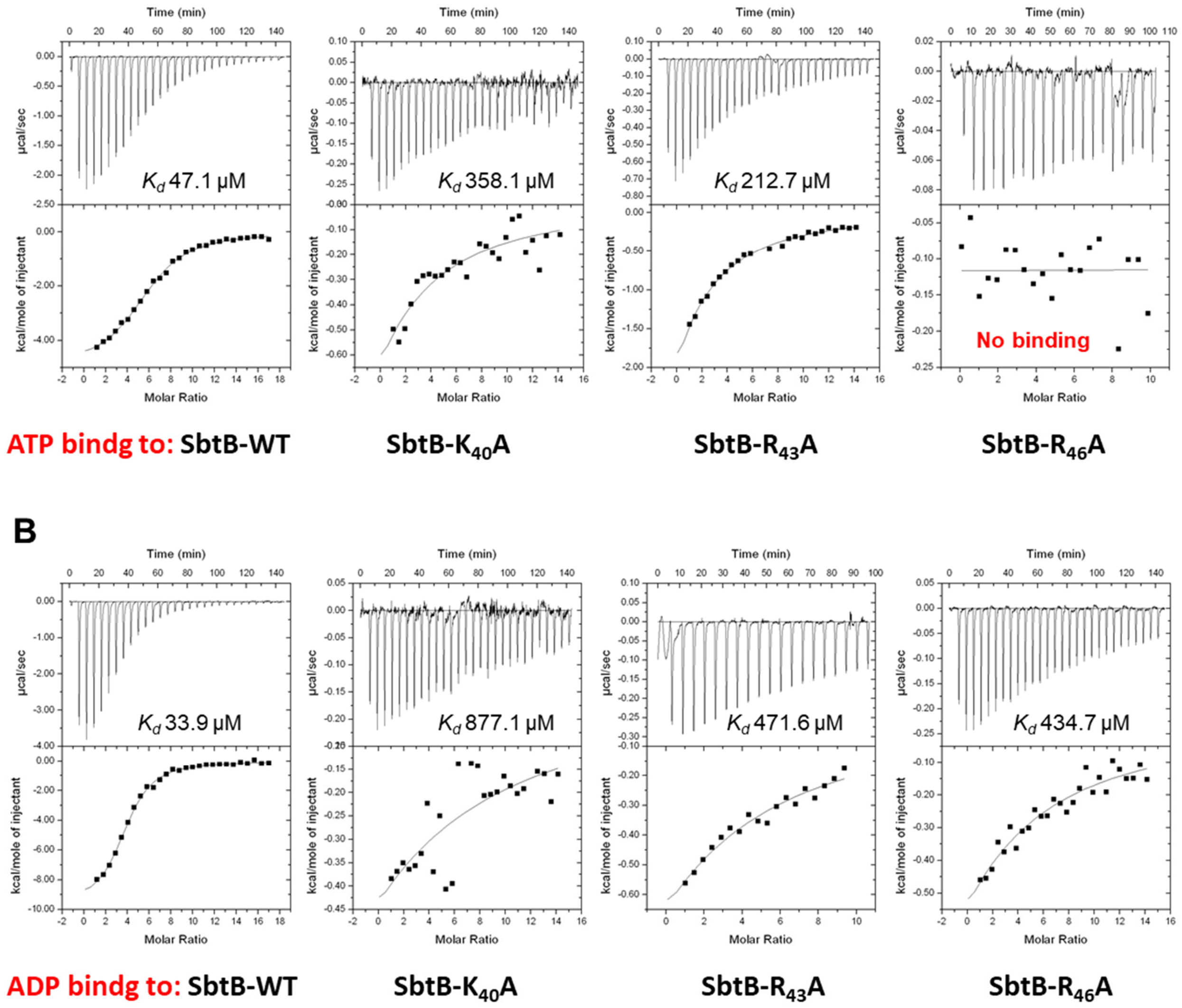


**Fig. S1: ITC analysis of SbtB and its T-loop variants (SbtB-K$_{40}$A, SbtB-R$_{43}$A and SbtB-R$_{46}$A).** Upper panels show the raw ITC data in the form of the heat produced during the titration of 30 µM SbtB (trimeric concentration) with ATP (A) and ADP (B); while lower panels show the binding isotherms and the best-fit curves to calculate $K_d$ values according to the one binding site models.

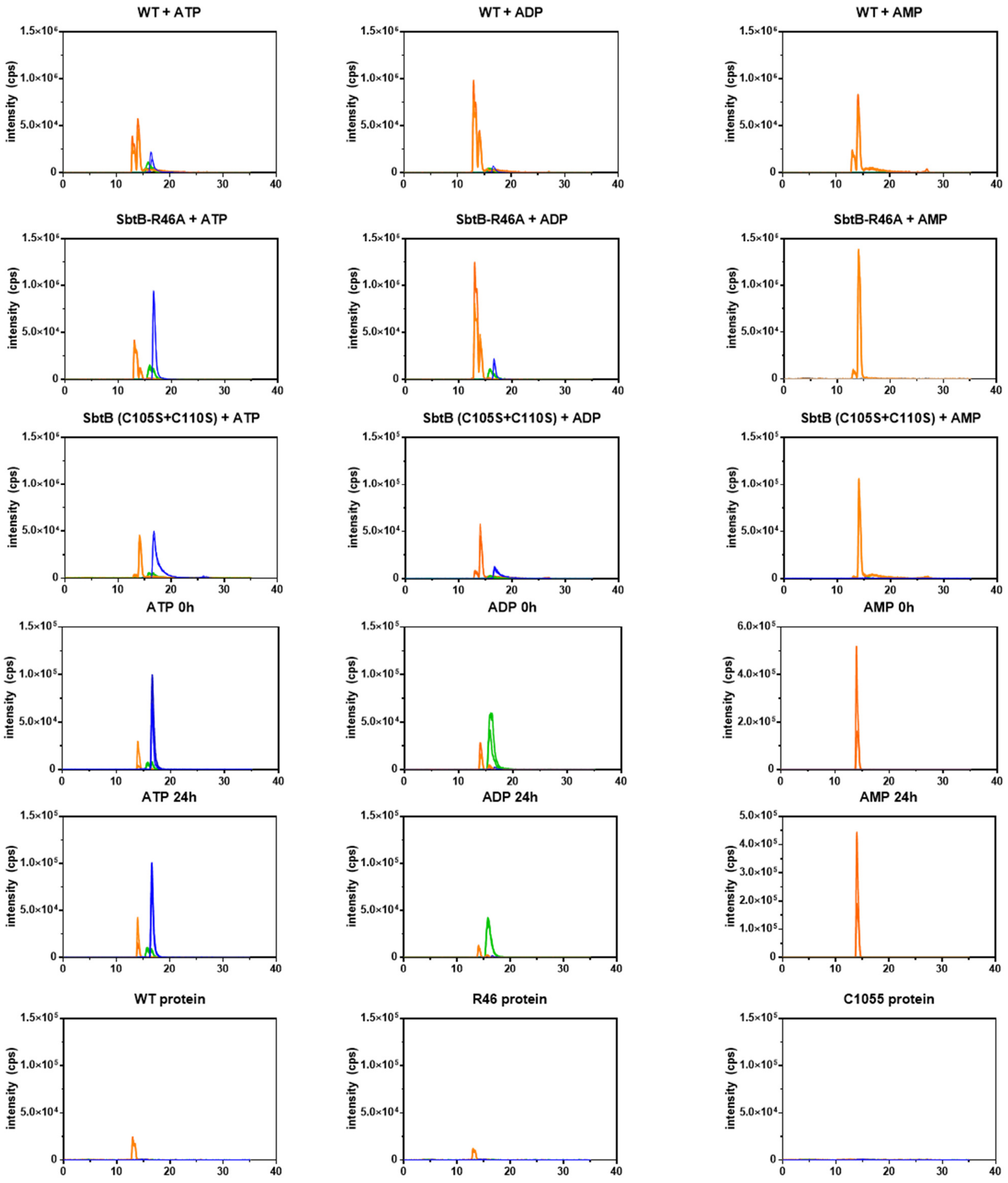


**Fig. S2:** HPLC-MS analysis was performed for SbtB wildtype and its variants ($R_{46}A$ and C105S+C110S [+ve control]) and the extracted ion chromatograms for ATP, ADP and AMP were generated, as indicated. The area under the curve for each nucleotide was used to quantify relative intensity of each nucleotide.

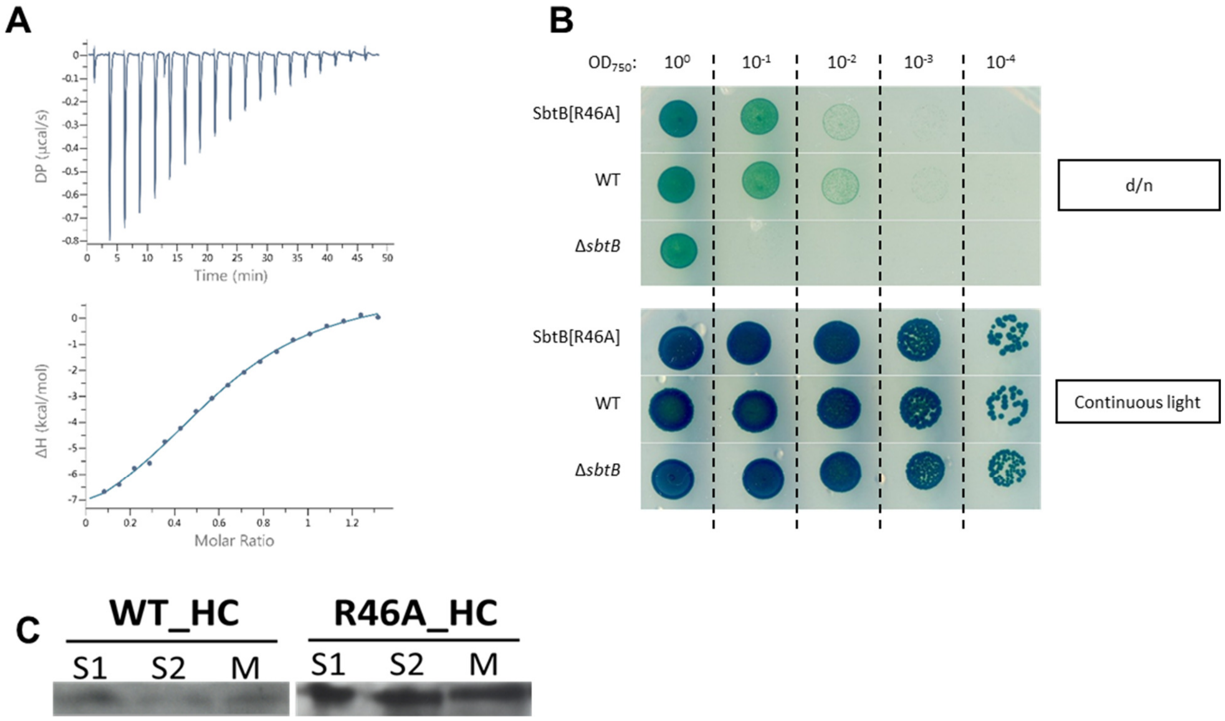


**Fig. S3: Characterization of SbtB-R$_{46}$A mutant.** (A) An isothermal calorimetric assay of recombinant SbtB-R$_{46}$A protein (37 µM; trimeric concentration) against 750 µM c-di-AMP. (B) While Δ*sbtB* had a severe growth defect in a diurnal rhythm, the *sbtB-R$_{46}$A* mutant did not show impaired growth compared to wildtype *Synechocystis* (upper picture). Cells were normalized to an OD$_{750}$ of 1.0 and serial diluted to an OD$_{750}$ of $1*10^{-4}$. Plates were either exposed to a diurnal rhythm (upper picture) or to continuous light (lower picture). (C) Analysis of SbtB localization. Localization of SbtB in soluble (S1 & S2) and membrane (M) fractions of cells grown under high carbon (HC; 5% $CO_2$) as assessed by immunoblot using anti-SbtB antibodies.

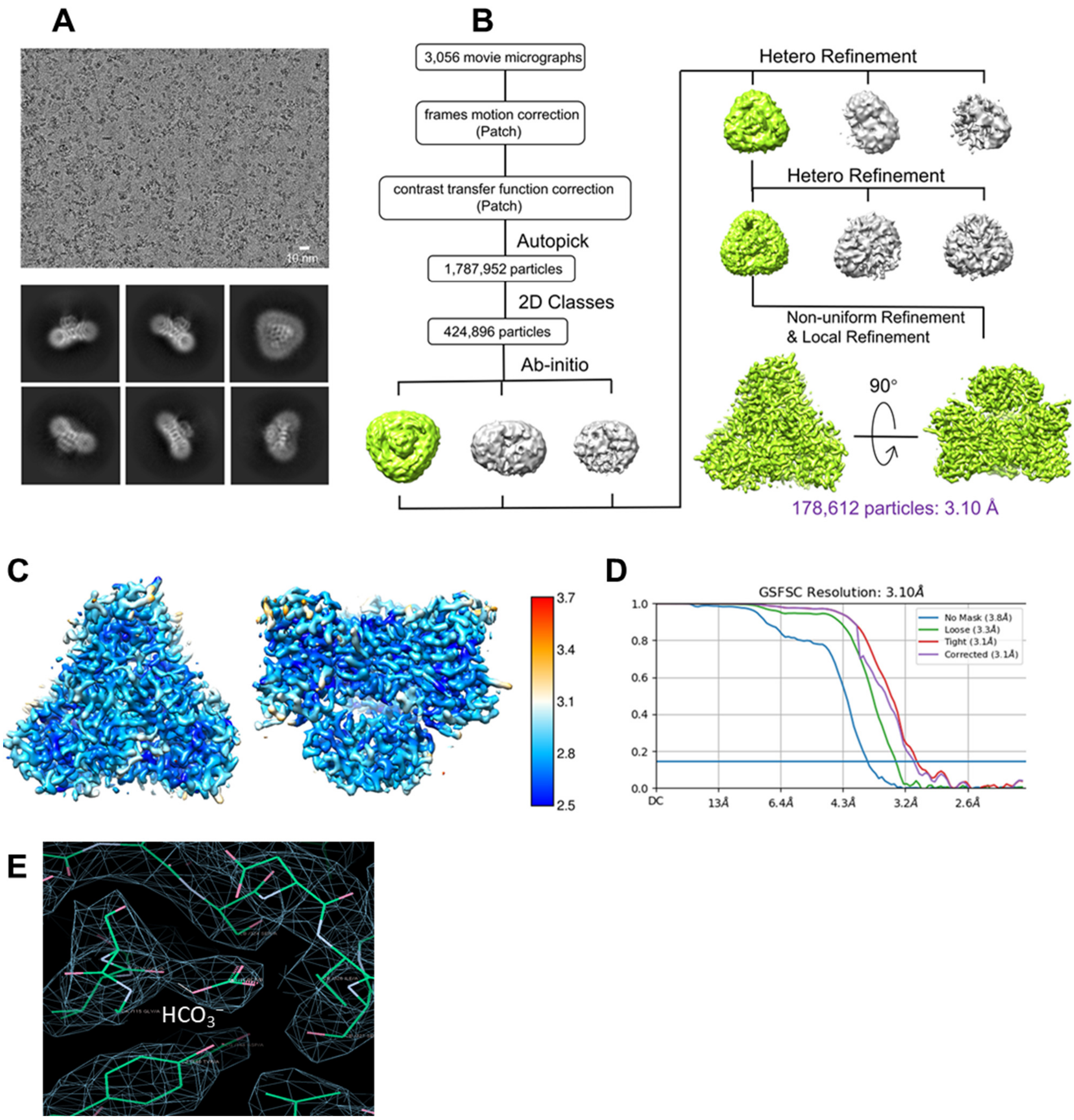


**Fig. S4: Cryo-EM analysis of SbtA:SbtB ΔT-loop complex of *Synechocystis* sp. PCC 6803.** (A) Representative cryo-EM micrograph of SbtA:SbtB ΔT-loop complex and 2D classes. (B) Data processing flowchart with particle distributions. (C) Resmap resolution slice and resolution map for SbtA:SbtB ΔT-loop complex, shown in the top view and the side view espectively. (D) Fourier shell correlation (FSC) curves showing a resolution of 3.1 Å. (E) The electron density of $HCO_3^-$.

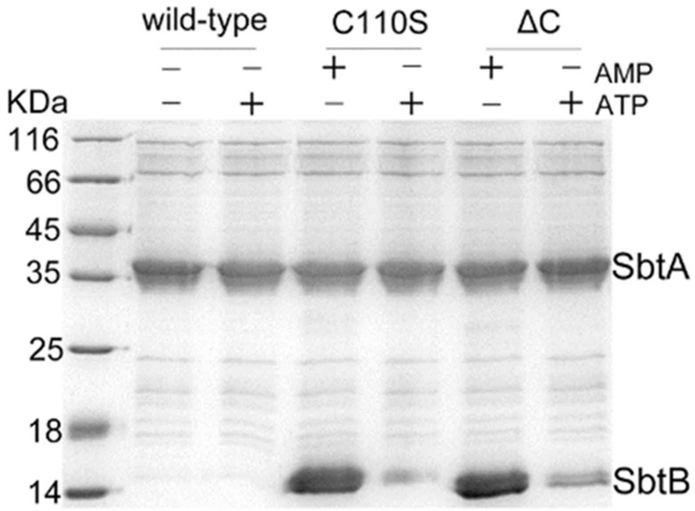


**Fig. S5:** Interaction of SbtA with SbtB and its variants (C105S and ΔC, in which the entire R-loop from $C_{105}$-$C_{110}$ was deleted) by pulldown assays in presence and absence of ATP or AMP as indicated.

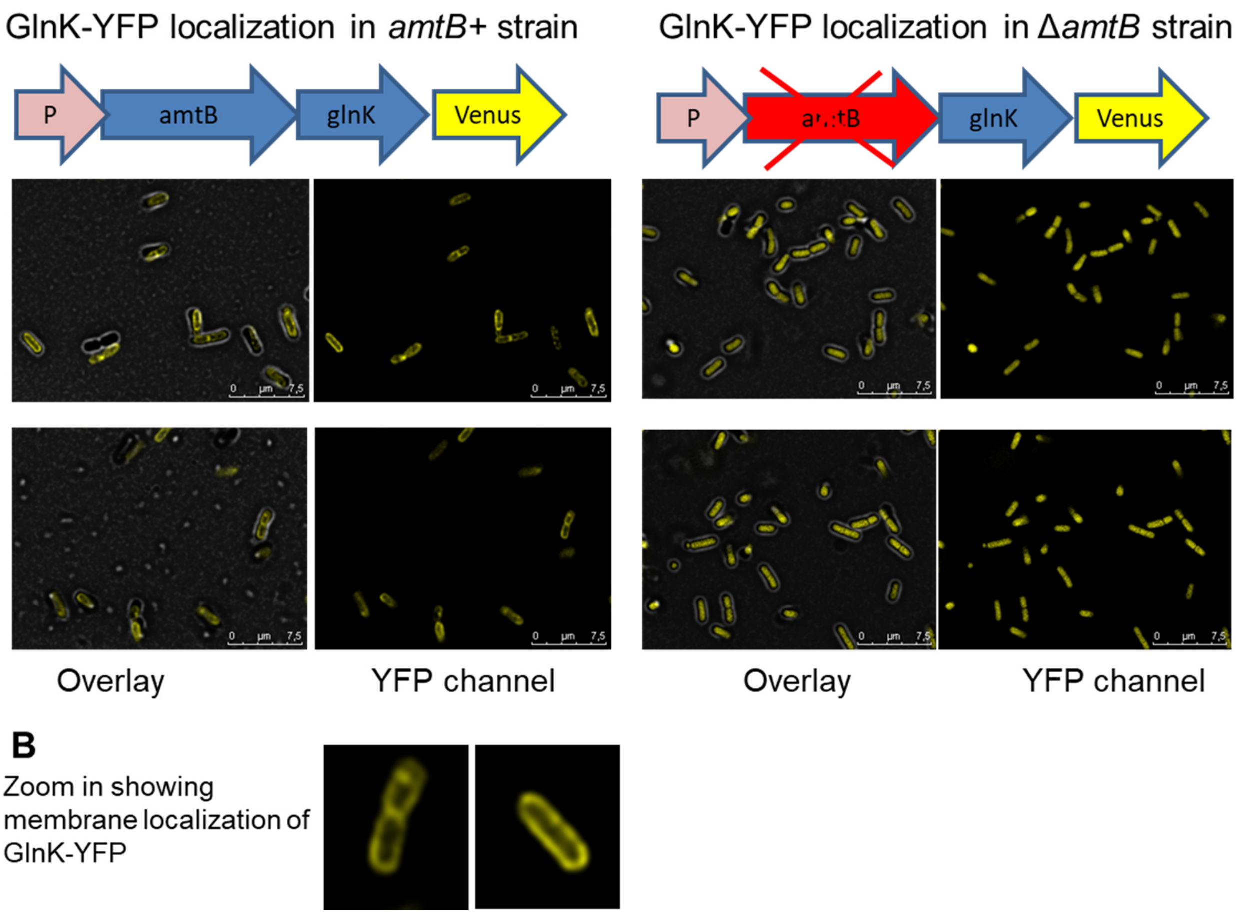


**Fig. S6:** GlnK localization in *amtB+* and Δ*amtB* strains under nitrate growth conditions as indicated.

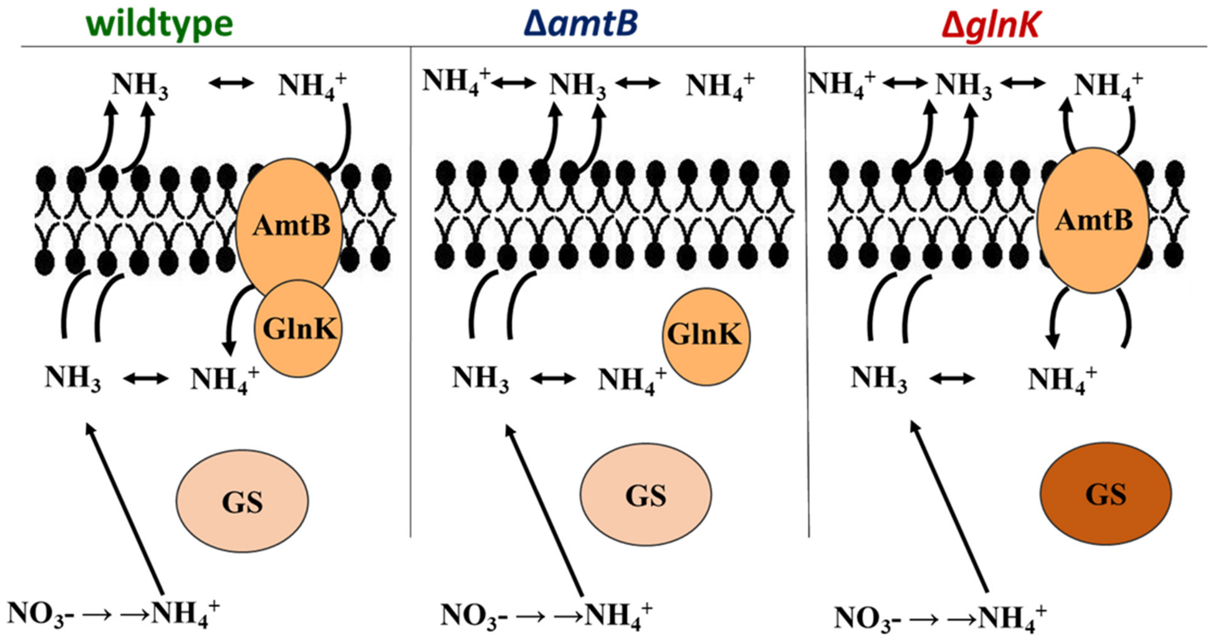


**Fig. S7:** Proposed mechanism for GlnK to act as a valve plug to control the ammonium transporter AmtB under poor nitrogen conditions in *Bacillus*.

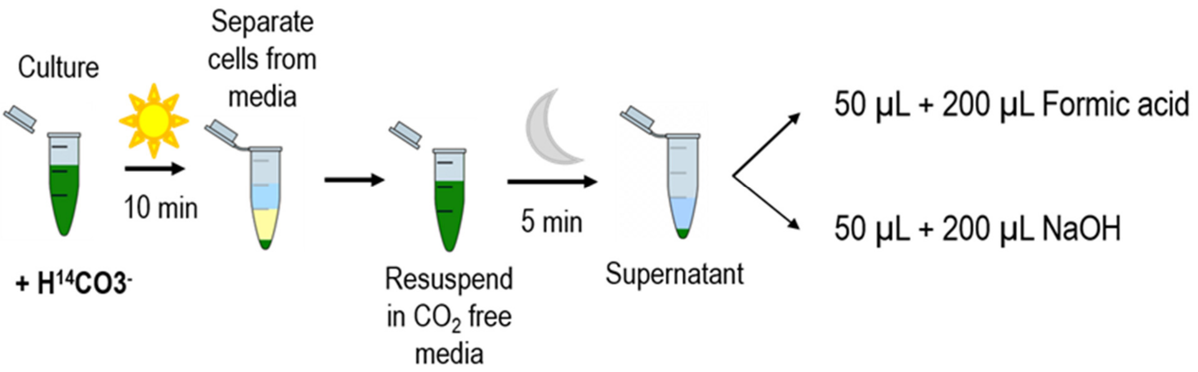


**Fig. S8:** Simplified scheme of $HCO_3^-$ leak experiment protocol.

### Supplementary Table S1. Primers and Plasmids List

| Primers/ amplification | Sequence (5'→3') | Note/ Ref. |
|---|---|---|
| **Creation of Δ*sbtB-R46A* mutant (pUC19-SbtB-$R_{46}$A)** | | |
| Sequencing primers for SbtB-R46A | pUC19-Seq-Fw: GCTGCAAGGCGATTAAGTTG | This study |
| | 116-RBSeq-Rv: ACACAGGAAACAGCTATGAC | This study |
| **Recombinant proteins** | | |
| WT *Sc*SbtB (*slr1513*) | 1256_Fw: GTGAAATGAATAGTTCGACAAAAATCTAGATAACGAGGGCAAAAAATG GCTAAACCAGCGAACAAGCTCG | Selim et al. 2018 |
| | 1257_Rv: AAGCTTATTATTTTTCGAACTGCGGGTGGCTCCAAGCGCTACAGCCCT CAGGGCCACAGAAAG | Selim et al. 2018 |
| *Sc*SbtB-ΔC | 1256_Fw: GTGAAATGAATAGTTCGACAAAAATCTAGATAACGAGGGCAAAAAATG GCTAAACCAGCGAACAAGCTCG | Selim et al. 2023 |
| | 1663_Rv_SbtB delta 104: CAAGCTTATTATTTTTCGAACTGCGGGTGGCTCCAAGCGCTGAAAGTATGCCCATAAAGTACTTCTGC | Selim et al. 2023 |
| *Sc*SbtB-$C_{105}$S+$C_{110}$S | 1256_Fw: GTGAAATGAATAGTTCGACAAAAATCTAGATAACGAGGGCAAAAAATG GCTAAACCAGCGAACAAGCTCG | Selim et al. 2023 |
| | 1761_Rv_SbtB-C105+110S CAAGCTTATTATTTTTCGAACTGCGGGTGGCTCCAAGCGCTGCTGCCCTCAGGGCCGCTGAAAGTATGC CCATAAAGTACTTCTGC | Selim et al. 2023 |
| *Sc*SbtB-$R_{46}$A | 1855_Fw_ R46A_SbtB: AATACCGGTGGCAAGGGTAGCCGTAACGTGGCCTCGTCGGGTCAAC | Selim et al. 2023 |
| | 1856_Rv_K40,R43,R46:A_SbtB: CATTACCGTGTATCCTTTGGCACCGGATTCG | Selim et al. 2023 |
| *Sc*SbtB-$R_{43}$A | 1854_Fw_R43A_SbtB: AATACCGGTGGCAAGGGTAGCGCCAACGTGCGCTCG | Selim et al. 2023 |
| | 1856_Rv_K40,R43,R46:A_SbtB: CATTACCGTGTATCCTTTGGCACCGGATTCG | Selim et al. 2023 |
| *Sc*SbtB-$K_{40}$A | 1853_Fw_K40A_SbtB: AATACCGGTGGCGCTGGTAGCCGTAAC | Selim et al. 2023 |
| | 1856_Rv_K40,R43,R46:A_SbtB: CATTACCGTGTATCCTTTGGCACCGGATTCG | Selim et al. 2023 |

**Supplementary Table 2.** Cryo-EM parameters, data collection and refinement statistics.

| | SbtA:SbtB ΔT-loop complex |
|---|---|
| **Data collection and processing** | |
| Magnification | 81,000 |
| Voltage (kV) | 300 |
| Electron exposure (e – /Å$^2$) | 55 |
| Defocus range (μm) | -1.2 to -2.0 |
| Pixel size (Å) | 1.07 |
| Symmetry imposed | C3 |
| Initial particle images (no.) | 1,787,982 |
| Final particle images (no.) | 424,896 |
| Map resolution (Å) | 3.10 |
| FSC threshold | 0.143 |
| Map resolution range (Å) | 2.02-999 |
| **Refinement** | |
| Initial model used (PDB code) | XXXX |
| Map sharpening B factor (Å$^2$) | -134.001 |
| Model resolution (Å) | 3.10 |
| FSC threshold | 0.143 |
| **Model composition** | |
| Non-hydrogen atoms | 9,402 |
| Protein residues | 1,452 |
| Waters | 0 |
| **R.M.S. deviations** | |
| Bond lengths (Å) | 0.009 |
| Bond angles (°) | 1.180 |
| **Validation** | |
| MolProbity score | 2.77 |
| Clashscore | 35.68 |
| Poor rotamers (%) | 2.85 |
| **Ramachandran plot** | |
| Favored (%) | 94.19 |
| Allowed (%) | 5.49 |
| Disallowed (%) | 0.32 |
